# Physiological roles of short-chain and long-chain menaquinones (vitamin K2) in *Lactococcus cremoris*

**DOI:** 10.1101/2021.11.27.470198

**Authors:** Yue Liu, Nikolaos Charamis, Sjef Boeren, Joost Blok, Alisha Geraldine Lewis, Eddy J. Smid, Tjakko Abee

## Abstract

*Lactococcus cremoris* and *L. lactis* are well-known for their occurrence and applications in dairy fermentations, but their niche extends to a range of natural and food production environments. *L. cremoris* and *L. lactis* produce MKs (vitamin K2), mainly as the long-chain forms represented by MK-9 and MK-8, and a detectable amount of short-chain forms represented by MK-3. The physiological significance of the different MK forms in the lifestyle of these bacterial species has not been investigated extensively. In this study, we used *L. cremoris* MG1363 to construct mutants producing different MK profiles by deletion of genes encoding (i) a menaquinone-specific isochorismate synthase, (ii) a geranyltranstransferase and (iii) a prenyl diphosphate synthase. These gene deletions resulted in (i) a non-MK producer (Δ*menF*), (ii) a presumed MK-1 producer (Δ*ispA*) and (iii) a MK-3 producer (Δ*llmg_0196*), respectively. By examining the phenotypes of the MG1363 wildtype strain and respective mutants, including biomass accumulation, stationary phase survival, oxygen consumption, primary metabolites, azo dye/copper reduction, and proteomes, under aerobic, anaerobic and respiration-permissive conditions, we could infer that short-chain MKs like MK-1 and MK-3 are preferred to mediate extracellular electron transfer and reaction with extracellular oxygen, while the long-chain MKs like MK-9 and MK-8 are more efficient in aerobic respiratory electron transport chain. The different electron transfer routes mediated by short-chain and long-chain MKs likely support growth and survival of *L. cremoris* in a range of (transiently) anaerobic and aerobic niches including food fermentations, highlighting the physiological significance of diverse MKs in *L. cremoris*.

## Introduction

*Lactococcus cremoris* [previous known as *Lactococcus lactis* ssp. *cremoris* (Li et al., 2021)] and *L. lactis* are lactic acid bacterium (LAB) that play important roles in food fermentation processes, especially in the manufacturing of fermented dairy products: they are the main constituent of various dairy starter cultures used all over the world for the production of cheese, butter milk and sour cream (Cavanagh et al., 2015). The essential involvement of *L. cremoris* and *L. lactis* in the fermentation of food raw materials highlights the interest in understanding the physiology and lifestyle of this bacterium.

Although *L. cremoris* and *L. lactis* are best known for their application and occurrence in dairy products, they are found in a diverse range of natural niches such as the gastrointestinal tract of particular fish species and various plant materials (Cavanagh et al., 2015). In fact, the dairy isolates are believed to originate from plant isolates which successful adapted to thrive in the dairy environment (Kleerebezem et al., 2020). It has been suggested that the niche adaptation history is reflected in the lifestyle and adaptability of *L. cremoris* and *L. lactis* in different environmental conditions. For instance, although *L. cremoris* is classified as a facultative anaerobe with a fermentative metabolism, evidence has also been provided that in presence of oxygen and exogenous supplemented of heme, the organism can switch to aerobic respiration, a process enabled by the menaquinones (MKs, also referred to as vitamin K2) produced by *L. cremoris* as electron carriers (Duwat et al., 2001; Brooijmans et al., 2009c). Notably, many other LAB species commonly applied in food fermentation processes, e.g., *Lactiplantibacillus plantarum* [previously referred to as *Lactobacillus plantarum* (Zheng et al., 2020)], have lost the ability to produce MKs (Pedersen et al., 2012). The ecological and physiological significance of MKs and their contribution to the successful applications of *L. cremoris* in food fermentations is an interesting, but under-explored topic.

MKs accumulate in the cell membrane of producing bacteria (Walther et al., 2013). All MKs share a naphthoquinone structure but the variants differ in the length of sidechains consisting of isoprenyl units (Lenaz and Genova, 2013). In the abbreviation MK-n, the number of isoprenyl units in the side chain is indicated by n. The variants of MKs produced in different bacterial species are distinct and have even been proposed as taxonomic markers (Collins and Jones, 1981). For example, *Bacillus subtilis* produces MK-7, *Escherichia coli* produces mainly MK-8 and *Propionibacterium freudenreichii* produces MK-9(4H) (Walther et al., 2013). *L. cremoris* and *L. lactis* are known to produce a range of MK variants, including MK-3, MK-5 through MK-10, among which MK-8 and MK-9 are the major forms (Brooijmans et al., 2009c; Liu et al., 2019).

MK is the sole quinone that shuttles electrons in the respiratory electron transport chain (ETC) in Gram-positive bacteria, essential for both aerobic and anaerobic respiration processes (Kurosu and Begari, 2010). For respiring Gram-positive bacteria, MK is essential for growth and survival, and enzymes in the biosynthesis pathway in pathogens have therefore been considered as targets for developing antibacterial agents. As mentioned, functional respiration in response to oxygen and heme supplementation has been observed *L. cremoris* strains, where the NADH dehydrogenase complex, MKs and the bd-type cytochrome complex (where heme is required as a cofactor) together form a functional ETC in the cell membrane (Brooijmans et al., 2009c). *L. cremoris* is not able to synthesis heme, thus heme has to be supplied exogenously to allow aerobic respiration in *L. cremoris* (Pedersen et al., 2012). Nevertheless, heme is not considered to be present in dairy environment, and aerobic respiration is not a common or essential metabolic mode for *L. cremoris* in food fermentations. In fact, in the production processes of fermented dairy product, *L. cremoris* is considered to mainly encounter anaerobic conditions during the fermentation. However, *L. cremoris* may be exposed to oxygen during starter culture production and during early stages in cheese production, i.e., aerobic conditions in absence of heme (Cretenet et al., 2014).

Under anaerobic conditions, MKs are known to endow some bacteria with the ability to utilize extracellular electron acceptors as alternative for oxygen, e.g., nitrate or fumarate, allowing anaerobic respiration as reported for *E. coli, Enterococcus faecalis and La. plantarum* (with exogenously supplied heme and MK). (Huycke et al., 2001; Brooijmans et al., 2009b; Kurosu and Begari, 2010). Moreover, MKs or precursors such as demethylmenaquinones (DMKs) or naphthoquinones were shown to be crucial for extracellular electron transport (EET) in various Gram-positive bacteria species including *L. cremoris*, which has been shown to be able to reduce Cu^2+^ ions, transfer electrons to electrodes and decolorize azo dyes conceivably via flavin-based EET (Rezaïki et al., 2008; Freguia et al., 2009; Pérez-Díaz and McFeeters, 2009; Light et al., 2018, 2019).

Under aerobic conditions, *L. cremoris* is known to be oxygen tolerant. (D)MKs were found to mediate reduction of exogenous oxygen to reactive oxygen species (ROS) such as superoxide in *L. cremoris* and *En. faecalis* (Huycke et al., 2001; Rezaïki et al., 2008). On the other hand, Liu et al. obtained *L. cremoris* vitamin K2 (MK) overproducers that showed high resistance to oxidative stress by laboratory evolution under aerobic conditions, although a direct relationship between elevated MK content and enhanced resistance to oxidative stress remains to be established (Liu et al., 2021).

Interestingly, a significant amount of MKs was found in *L. cremoris* stationary phase cells from different growth conditions: anaerobic/microaerophilic, aerobic and respiration-permissive (aerobic conditions with addition of heme) (Brooijmans et al., 2009c; Liu et al., 2019). Notably, *L. cremoris* produces multiple MK forms, whereas most MK producing bacteria are known to produce one specific form, or one major form with two adjacent forms in very minor amount. *L. cremoris* produces noticeable amounts of both short-chain MKs represented by MK-3, and long-chain MKs represented by MK-8 and MK-9, and the distribution of the short-chain and long-chain MK forms alters depending on the growth conditions (Brooijmans et al., 2009c; Walther et al., 2013; Liu et al., 2019). It remains to be elucidated whether the short-chain or long-chain forms of MKs are favored for particular functions under the different aeration conditions (anaerobic or aerobic) and metabolic modes (fermentation or respiration) that *L. cremoris* could be exposed to in a range of natural and food production environments, for example in dairy fermentations.

To examine the physiological significance of the short-chain and long-chain MKs in *L. cremoris*, we constructed dedicated mutants using model strain *L. cremoris* MG1363, based on annotation of putative genes in the MK biosynthesis pathway in *L. cremoris* (Fig. 1) (Wegmann et al., 2007)(Brooijmans et al., 2009c): candidate genes encoding a menaquinone-specific isochorismate synthase (*menF*), a geranyltranstransferase (*ispA*) and a prenyl diphosphate synthase (*llmg_0196*) were deleted, with the prediction to create mutants producing no MK (Δ*menF*), only MK-1 (Δ*ispA*) and only MK-3 (Δ*llmg_0196*) respectively. These mutants, together with the original strain MG1363 producing mainly MK-9 and MK-8, allowed the study of functionality of short-chain and long-chain MK forms in *L. cremoris* under anaerobic, aerobic and respiration-permissive conditions, where involvement of MKs in bacterial metabolism and physiology has been proposed. Under these conditions, relevant phenotypes including MK profile, biomass accumulation, viability/survival, oxygen consumption, electron transfer to extracellular acceptors (e.g., azo dye, Cu^2+^), metabolites and proteomes were examined for strain MG1363 and mutants.

**Figure 1.**
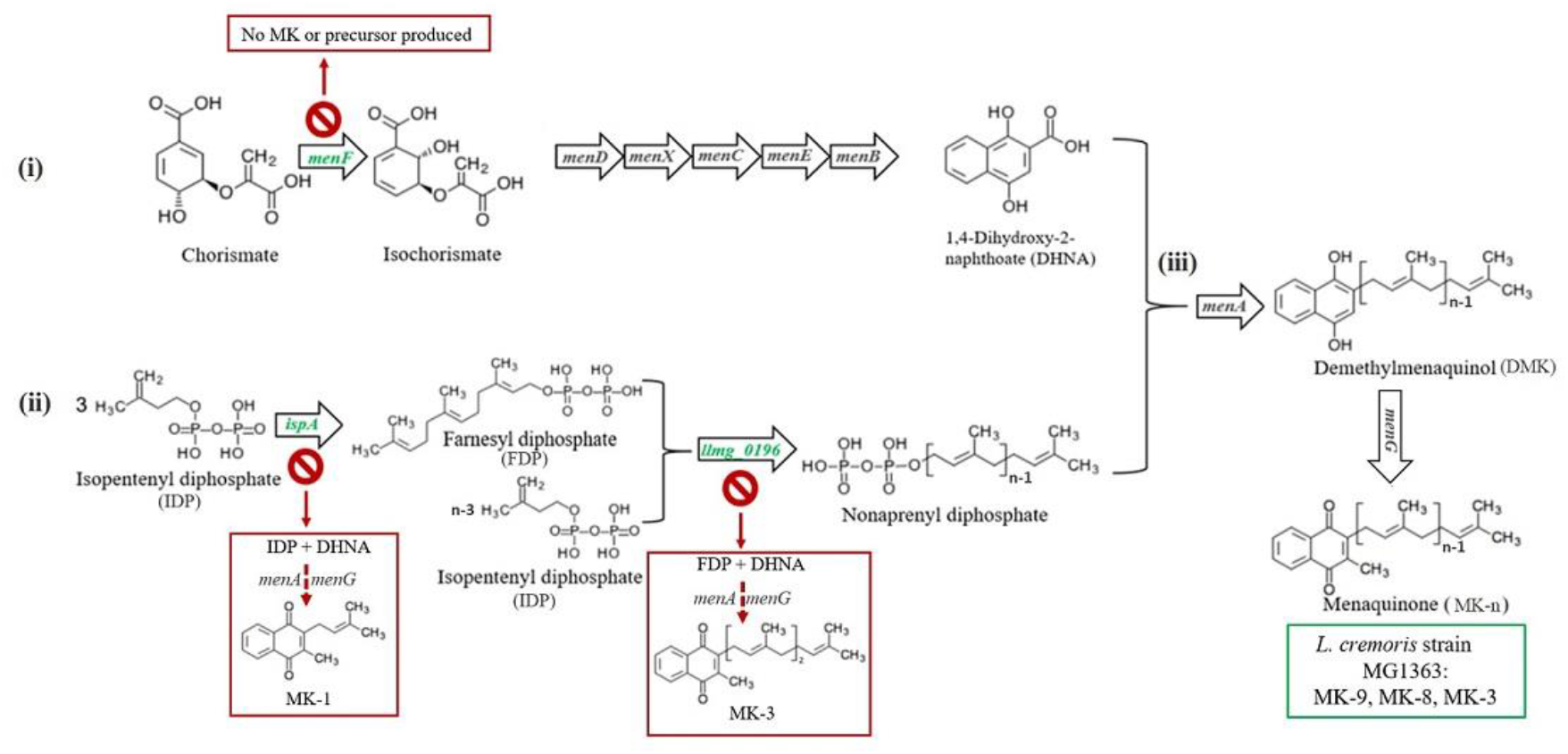
Predicted menaquinone biosynthesis pathway in *L. cremoris* strain MG1363 and mutants. MK biosynthesis in *L. cremoris*/*L. lactis* is predicted to follow a pathway consisting of three parts: (i) synthesis of the naphthoquinone structure (the head group), (ii) synthesis of the isoprenyl side chain (the tail group), and (iii) joining of the head and tail structures giving the MK. The head group synthesis starts with chorismate generated from the Shikimate pathway with the first step catalyzed by a menaquinone-specific isochorismate synthase (predicted encoding gene is *menF*), and the tail group synthesis starts with isopentenyl diphosphate (IPP), containing a single isoprene unit, generated from mevalonate pathway. For building the side chain, first a geranyltranstransferase (predicted encoding gene is *ispA*) is required to form farnesyl diphosphate (FPP) containing three isoprene units. More IPP is linked to FPP to form longer isoprene chains, and this step is catalyzed by a prenyl diphosphate synthase (predicted encoding gene is *llmg_0196*). The latter enzyme also is thought to play a role in determination of the length of the isoprene chain (Nagel et al., 2018; Johnston and Bulloch, 2020). Finally, the naphthoquinone head group - 1.4-dihydroxy-2-naphthoic acid (DHNA), and the polyprenyl tail group are combined by isoprenyl diphosphate:1,4-dihydroxy-2-naphthoate isoprenyltransferase to form demethylmenaquinone (DMK), which is methylated to give MK. Mainly MK-9, MK-8, MK-3 are formed in original *L. cremoris* strain MG1363 (green box). Mutants Δ*menF*, Δ*ispA and* Δ*llmg_0196* are expected to produce no MK, only MK-1 and only MK-3, respectively. Genes targeted in this study shown in green color. Predicted MK production route and profiles in respective gene deletion mutants are shown in red boxes.

## Materials and methods

### Strains and conditions

*Lactococcus cremoris* MG1363 and derived gene deletion mutants Δ*llmg_0196*, Δ*ispA* and Δ*menF* were cultivated in GM17 medium [M17 broth (Difco, BD Biosciences) supplemented with 0.5 % (w/v) glucose] at 30 °C. Cultivation times and conditions are specified in the Results section and figure legends for each particular case. For mutants with gene complementation, 5 μg/mL erythromycin was added to the medium.

For general static cultivation, cultures were incubated statically in closed Greiner tubes in maximal volume under normal atmosphere. For specified anaerobic cultivation, cultivation vessels with bacterial cultures were placed in anaerobic jars flushed with gas mix (10% CO_2_, 10% H_2_ and 80% N_2_) using the anaerobic cycle programme by Anoxomat (WS9000, Mart Microbiology, Netherlands) unless specified otherwise. For aerobic cultivation, bacterial cultures filled up to 10% volume of Erlenmeyer flasks and were shaken at 180 rpm. Respiration-permissive cultivation conditions were the same as those applied for aerobic cultivations but with the addition of 2 μg/mL heme (hemin, Sigma).

*E. coli* strains used in the genetic modification procedures, harbouring plasmids with an erythromycin resistance gene (*ermAM*), were cultivated/selected in LB medium supplemented with 150 μg/mL erythromycin at 37 °C. Liquid cultures were shaken at 160 rpm with 90% headspace for no longer than 16 h, agar plates were incubated at normal atmosphere for maximal 22 h.

### Genetic modification

#### Selection of targeted genes

The MK biosynthesis pathway in *L. cremoris* MG1363 was predicted based on information retrieved from KEGG (Kanehisa and Goto, 2000) and genome of *L. cremoris* MG1363 (Wegmann et al., 2007) and homology searches based on sequences from other bacteria with experimentally confirmed functions (Fig. 1). To obtain mutants of *L. cremoris* MG1363 with different MK profiles, several genes were selected and subsequently targeted for gene deletion procedures. For the mutant producing no MK at all, it was decided that the gene involved in the first step of MK-specific synthesis pathway would be the target, to eliminate possible interference of intermediate products on the mutant phenotype (Fig. 1). We used BLAST to identify MenF homologs in the microbial protein database (Joint Genome Institute) (Chen et al., 2019). Gene product of *menF* (*llmg_1828*) in *L. cremoris* MG1363 was identified with 26% identity with MenF protein sequence (Uniprot ID: P38051) from *E*.*coli* K12, for which the function has been confirmed (Daruwala et al., 1996), and was chosen as the first target gene to delete in strain MG1363. For the mutant producing short-chain MKs, genes encoding the geranyltranstransferase and prenyl diphosphate synthase were the targets. Combining previous knowledge (Kobayashi et al., 2003), sequences of genes *hepT* (P31114), *hepS* (P31112) and *yqiD* (P54383) from *Bacillus subtilis* 168 were used to search for homologs in *L. cremoris* MG1363 using BLAST. No sequences similar to *B. subtilis hepS* were found in *L. cremoris* MG1363. Moreover, *ispB* (*llmg_1110*), *llmg_0196* and *ispA* (*llmg_1689*) in *L. cremoris* MG1363 were identified with 34%, 31% and 35% identity to *B. subtilis hepT* respectively. Finally, *ispB* (*llmg_1110*), *llmg_0196* and *ispA* (*llmg_1689*) were identified with 30%, 28% and 45% identity to *B. subtilis yqiD* respectively. Genes encoding *ispB* (*llmg_1110*), *llmg_0196* and *ispA* (*llmg_1689*) were all chosen as the target genes for modifying MK side chains to obtain mutants producing only MK-3 or MK-1 (Fig. 1). The *ispB* (*llmg_1110*) deletion mutant did not show any alteration in MK profile compared to MG1363 and will not be discussed in further details in this study.

#### Plasmid construction

Gene deletion in *L. cremoris* MG1363 was achieved by homologous recombination. To construct the plasmids for gene deletion, the upstream and downstream homologous regions of 700 - 800 bp of the target genes were amplified, and desired restriction sites were introduced by PCR using primers listed in supplementary Table S1. Phusion High-fidelity PCR kit (Thermo Fisher Scientific) was used according to manufacturer’s instruction. Both upstream and downstream homologous regions of each target gene were inserted in plasmid pCS1966 [gift from Solem et al. (Solem et al., 2008)] by restriction digestion and ligation following basic guidelines provided with the enzymes (Thermo Fisher Scientific). In brief, restriction site PstI was used to connect the upstream and downstream homologous regions of each target gene, HindIII and XbaI sites were used to insert the two homologous regions to the backbone of pCS1966. As a result, plasmid pYL005, pYL004 and pYL003 harboring the surrounding homologous regions of *menF, llmg_0196* and *ispA*, respectively, were constructed.

For gene complementation, promoter regions (ca. 50 bp upstream of the start codon) and open reading frames of *menF, llmg_0196* and *ispA* were amplified by PCR using primers listed in supplementary Table S2. PCR products for each gene were inserted in the backbone (8301 bp fragment obtained by XbaI and BgIII digestion) of plasmid pMSP3545 [gift from Gary Dunny (Bryan et al., 2000), Addgene plasmid # 46888] using NEBuilder HiFi DNA assembly cloning kit (New England Biolabs) according to manufacturer’s instruction. As a result, p45-*menF*, p45-0196 and p45-*ispA* containing the promoter and gene sequences of *menF, llmg_0196* and *ispA*, respectively, were constructed.

Assembled plasmids were first transformed into *E. coli* competent cells Mix & Go Zymo 5α (Zymo research) according to product manuals, and transformed candidates were made into liquid culture for plasmid isolation. Assembled plasmids were checked for correctness by restriction digestion analysis after propagation in transformed *E. coli*. The correctness of PCR amplified sequences were checked by Sanger sequencing (BaseClear, Leiden, Netherlands).

#### Transformation and homologous recombination in L. cremoris

For gene deletion, plasmid pYL005, pYL004 and pYL003 were separately transformed into MG1363 with the protocol described by (Holo and Nes, 1989) with the following modifications: MG1363 was cultured in SMGG media (M17, 0.5 M Sucrose, 0.5% Glucose and 0.5% Glycine) till mid-log phase (OD_600_ = 0.6 - 0.9). An additional washing step was performed in between the original two washing steps, with 0.5 volume EDTA washing buffer (0.05 M EDTA, 0.5 M sucrose, 10% glycerol). For transformation, 500 – 1000 ng plasmid pYL005, pYL004 or pYL003 in volume 1 - 2 μL was added. Electroporation was done using the Gene Pulser Xcell Electroporation Systems (Bio-Rad) at 2500 V, 25 μF, 200 Ω. Cells were plated on selection plates (GM17, 1.5% agar, 0.5 M sucrose and 3 μg/mL erythromycin), and were incubated at 30 °C for 2-3 days till colonies emerge.

At this stage the plasmids, having no replication origin in *L. cremoris*, are integrated into the genome of MG1363 at either homologous region of the target genes. To eliminate the plasmid backbone and achieve gene deletion, transformants were inoculated in 2 mL SA medium (Jensen and Hammer, 1993) containing 1% glucose and incubated at 30 °C overnight. Then they were diluted 10x in SA (1% glucose) medium and incubated at 30 °C for 6 h. The culture was then plated on SA (1% glucose) agar plates supplemented with 10 μg/mL 5-Fluoroorotate (Sigma). Plates were incubated at 30 °C until colonies emerge. Due to the *oroP* gene on the plasmid of pCS1966, the presence of 5-fluoroorotate selects for mutants that have eliminated the plasmid backbone. From these mutants, the ones with gene deletion were selected by PCR (DreamTaq DNA polymerase, Thermo Fisher Scientific) using the primers listed in supplementary material Table S3.

For gene complementation, p45-*menF*, p45-0196 and p45-*ispA* were transformed into mutants Δ*menF*, Δ*ispA* and *llmg_0196* respectively with the electroporation protocol described above. Transformants were checked by PCR (DreamTaq DNA polymerase, Thermo Fisher Scientific) using the primers listed in supplementary material Table S2.

### MK analysis

MKs were extracted from cells as described previously (Liu et al., 2019). Briefly, the biomass was first treated with lysozyme and then MKs were extracted with organic solvent n-hexane and eventually dissolved in iso-propanol. All samples were diluted in methanol and subjected to analysis in the ultra-performance liquid chromatography (Thermo Scientific Vanquish) coupled with mass spectrometry (Thermo Q-Exactive hybrid quadrupole-Orbitrap) (UPLC–MS), exactly as described in (Liu et al., 2021). Calculations from analytical standards were performed as described previously (Liu et al., 2019).

### Biomass quantification

For cell dry weight (CDW) determination, PBS washed cell pellets from bacterial cultures were kept at 80 °C for 48-72 h to evaporate water content. The dried biomass was then weighed.

For optical density (OD) determination, OD_600_ of cell cultures was measured by a spectrophotometer (600 nm; path length 10 mm).

### Viable cell count enumeration

Bacterial cultures were subjected to serial dilution series in 96-well plates using PBS. Each dilution was spotted (10 μL) in triplicate on GM17 agar plates. Agar plates were incubated at 30 °C for 24 hours under microaerobic conditions before colonies were counted.

### Oxygen consumption rate analysis

The measurement and calculation of oxygen consumption rate in *L. cremoris* were performed exactly as described previously (Liu et al., 2021).

### Decolorization of azo dyes

Most azo dyes are not permeable to the cell membrane, and get decolorized upon reduction (Fang et al., 2019). Therefore, here we use the decolorization of an azo dye as an indicator for EET.

Strains were all cultivated under anaerobic conditions overnight in GM17 to obtain biomass for oxygen consumption analysis. The cells were harvested and washed in PBS once, and OD_600_ was standardized to 1 in PBS. Azo dye Reactive Black 5 (Sigma) was added to the cell suspension at a final concentration of 0.005%. The infusion bottles were flushed with N_2_ gas for 2 min to ensure an anaerobic environment. Reaction was at 30 °C and was initiated by adding glucose to a final concentration of 1%. OD of cell-free supernatant was followed for 4 h with 1 h interval at 595 nm. The OD_595_ at time 0 h was regarded as 100% colour intensity. The absolute value of the initial slope of the linear correlation between colour intensity and time for each measurement was taken as the absolute azo dye decolorization rate. Thereafter, a relative azo dye reduction rate was calculated by comparing the absolute decolorization rate among the strains, setting the highest absolute decolorization rate as 100%.

### Reduction of copper ions

The reduction of Cu^2+^ was measured indirectly in *L. cremoris*. When Cu^2+^ is reduced to the toxic species Cu^+^, growth of *L. lactis* was shown to be inhibited (Abicht et al., 2013). Therefore, inhibition of growth of *L. cremoris* was used as an indication of Cu^2+^ reduction.

Strains were cultivated statically in GM17 overnight, and then the cultures were diluted to an OD_600_ value of 0.1 in fresh GM17. For Cu^2+^ reduction tests, 300 nM CuCl_2_ was added to the media, and for growth controls CuCl_2_ was not added. For anaerobic cultivation, aliquots of 450 μL were brought into 100-well Honeycomb microplates (ThermoFisher) and incubated at 30 °C for 48 h in Bioscreen C (ThermoFisher), where the OD_600_ was measured at 1 h intervals. For aerobic cultivation, aliquots of 350 μL were transferred to microplates which were subsequently incubated with continuous shaking at medium intensity. For respiration-permissive conditions, cultures were incubated the same way as for aerobic conditions, albeit with the addition of 2 μg/mL heme (hemin, Sigma).

The time to reach (TTR) OD_600_ value 0.4 was used as a measurement of growth; the factor difference between TTRs in the test of 300 nM CuCl_2_ and in the growth control without CuCl_2_ was eventually used as the indicator of Cu^2+^ reduction-induced growth inhibition/toxicity.

### Metabolite analysis

Metabolites including lactate, formate, acetate, acetoin, ethanol and succinate were measured in cell-free supernatant from overnight cultures by high-performance liquid chromatography (HPLC) analysis exactly as described previously (Liu et al., 2021). The amount of each metabolite measured in uncultured GM17 medium was used as level zero to calculate the production or consumption of metabolites in the bacterial cultures.

### Proteomics analysis

Cells from overnight culture (16 h) were used for proteomics analysis. For each strain and condition combination, samples were collected from three independent experiments. Sample preparation, LCMS analysis and data processing were performed exactly as described previously (Liu et al., 2021). The proteome of *L. cremoris* MG1363 (UniProt ID UP000000364) was used as the protein database.

### Data analysis

Statistical significance analysis was performed in JASP (0.11.1) (Love et al., 2019) using two-way analysis of variance (ANOVA). Post hoc multiple comparisons were conducted using Tukey’s test (2-sided) and in all cases the control group was *L. cremoris* MG1363 (*P ≤ 0.05).

## Results

### MK profiles in mutants Δ*menF*, Δ*ispA* and Δ*llmg_0196*

First, we evaluated the gene deletion mutants derived from *L. cremoris* strain MG1363, by examining whether the MK profiles of the respective mutants were as predicted *in silico*. Wildtype strain MG1363 is known to produce mainly MK-8 and MK-9, but also MK-3 and a minor amount of other forms. The three gene deletion mutants were predicted to produce either no MK (Δ*menF*), only MK-1 (Δ*ispA*) or only MK-3 (Δ*llmg_0196*) based on the putative roles of respective gene products in the MK biosynthesis pathway (Fig. 1). The MK profiles as well as other phenotypes of strain MG1363 and the mutants were evaluated under three cultivation conditions (Fig. 2A): anaerobic, aerobic and respiration-permissive (meaning aerobic cultivation with heme supplementation).

**Figure 2.**
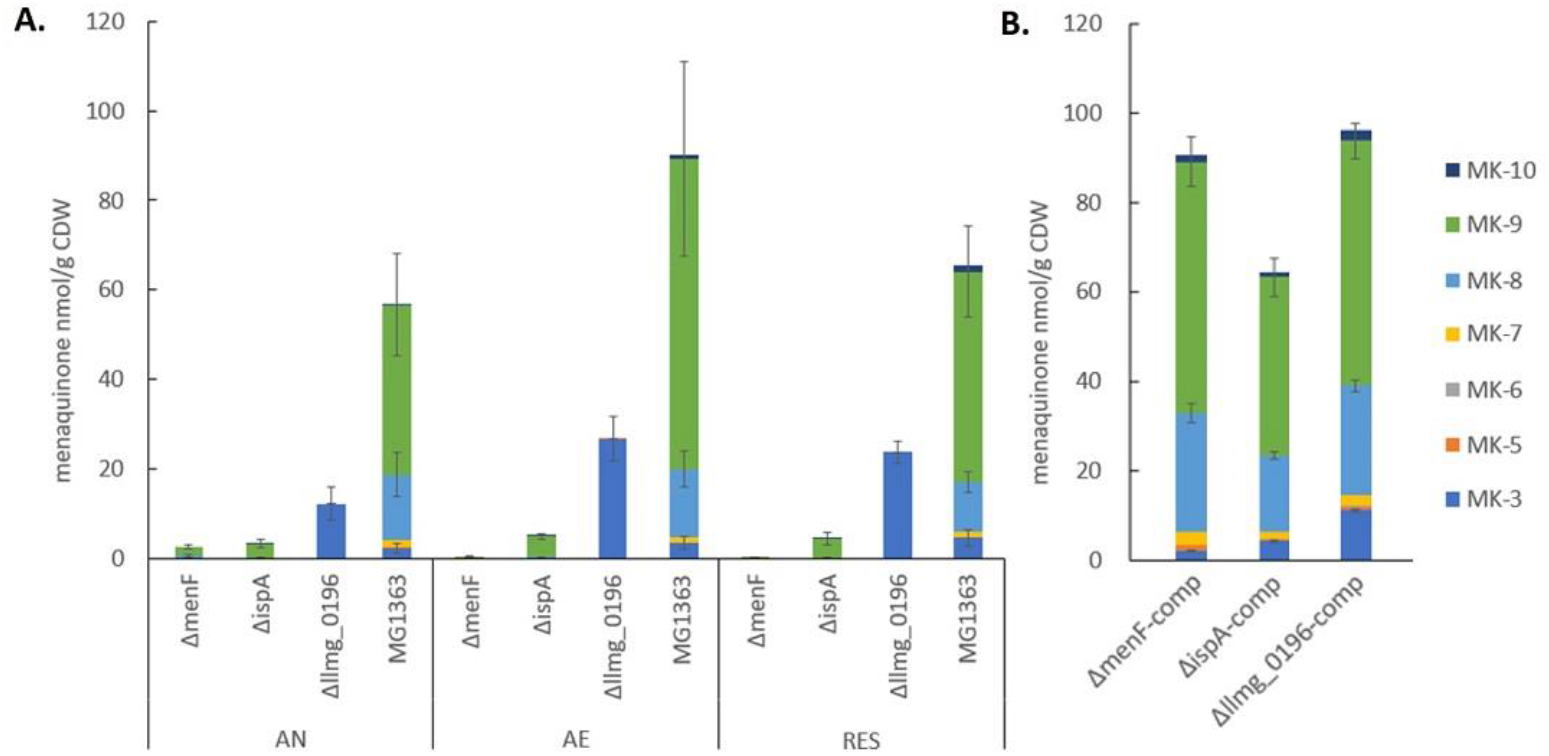
MK content in the biomass of *L. cremoris* strains. A) MK content in strain MG1363 and mutants with gene deletions. All strains were cultivated in GM17 media under indicated conditions for 16-18 hours at 30 °C before MKs were extracted from the biomass. AN: anaerobic; AE: aerobic; RES: respiration-permissive, i.e. supplement of both heme and aeration. Data from three independent experiments. B) MK content in mutants with respective gene complementation. Strains were cultivated under anaerobic conditions. Data from four biological replicates. Error bars show SEM for MK-3, MK-8 and MK-9.

Under all three tested cultivation conditions, substantial amounts of MKs mainly in the forms of MK-8 and MK-9, and noticeable amounts of MK-3 could be found in the biomass of wildtype strain MG1363. In addition, minor amounts of MK7 and MK-10 were also detected (Fig. 2A, supplementary Fig. S1A). The total amount of MKs in MG1363 was highest when cultivated under aerobic conditions, reaching 90 nmol/g cell dry weight (CDW), followed by respiration-permissive conditions of approximately 70 nmol/g CDW, while the amount found in cultures growing under anaerobic conditions was the lowest, less than 60 nmol/g CDW.

MG1363Δ*llmg_0196* produced solely MK-3 in the biomass, with the highest amount of about 25 nmol/g CDW was observed under aerobic and respiration-permissive conditions, and 12 nmol/g CDW under anaerobic conditions, confirming the hypothesis that gene *llmg_0196* encodes a (nona)prenyl diphosphate synthase. The molar amount of MK-3 produced in *Δllmg_0196* was higher than the MK-3 amount found in wildtype MG1363, but counted up to only one-third of the total MK amount in MG1363.

It was expected that mutant Δ*menF* would not produce MKs at all, and Δ*ispA* would produce only MK-1. Indeed, MG1363Δ*menF* showed no MK production except for trace amounts of MK-9 under anaerobic conditions (2 nmol/g CDW). MG1363Δ*ispA* showed low concentrations (3-5 nmol/g CDW) of MK-9 in all three conditions, while MK-1 was not detected, possibly due to limitations of the extraction method.

Besides examining the MK profiles in the biomass of *L. cremoris*, the cell-free supernatants of bacterial cultures were also examined for the presence of MKs. In all strains, no MK was detected in the culture supernatant except for Δ*llmg_0196* culture: a detectable amount of MK-3 was found in the supernatant, counting to 2% of the MK-3 quantity found in the cells obtained from the same culture.

When the mutants were complemented with the respective genes, the MK production levels were restored and profiles were similar to the original MG1363 (Fig. 2B, supplementary Fig. S1B).

### Anaerobic conditions: mutants showed distinct phenotypes in azo dye and Cu^2+^ reduction

Under anaerobic conditions, all mutants showed similar biomass accumulation as compared to wildtype MG1363 after overnight cultivation, which was about 1 g/L in cell dry weight (Fig. 3A, and OD measurement in supplementary Fig. S2A showed identical trend). The presumed MK-1 producer *ΔispA* and MK-3 producer *Δllmg_0196* showed 1-2 log higher survival than non-MK producer *ΔmenF* and wildtype MG1363 (MK-9, 8, 3 producer) after prolonged cultivation for 72 h (Fig. 3C).

**Figure 3.**
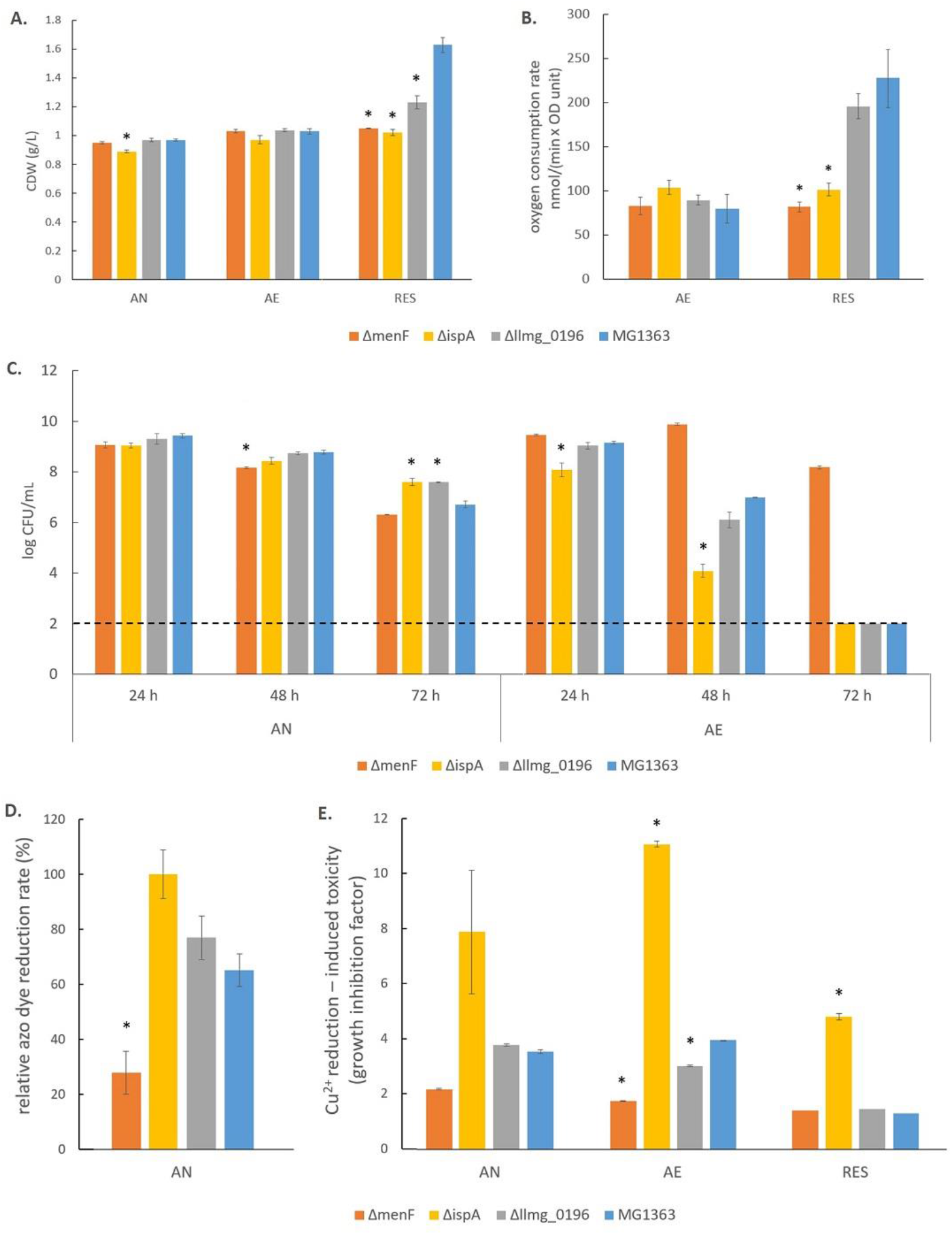
Phenotypical characterization of *L. cremoris* MG1363 and mutants under anaerobic, aerobic and respiration-permissive conditions. A) Biomass accumulation. Cell dry weight (CDW) was measured for strains cultivated in GM17 medium under respective conditions for 16-18 hours at 30 °C. B) Oxygen consumption rate. For AE test, cells were obtained from overnight culture in GM17 media under anaerobic conditions. For RES test, cells were obtained from overnight culture in GM17 media supplemented with heme under anaerobic conditions. PBS-washed cells were suspended in air-saturated PBS (OD standardize all to 2) for oxygen consumption test at room temperature. Reaction was initiated by adding 1% glucose. C) Viable plate count. All strains were cultivated in GM17 media under indicated conditions at 30 °C, culturability was determined at 24 h, 48 h and 72 h. All strains were inoculated at 10^6^ CFU/mL at 0 h. The dotted line indicates the detection limit. D) Relative azo dye reduction rate. Biomass was obtained from overnight culture under anaerobic conditions, washed and suspended in PBS (OD standardized to 1). Azo dye Reactive Black 5 was added to 0.005%. Reaction was at 30 °C and was initiated by adding 1% glucose. Initial azo dye reduction rate was measured by monitoring the decrease in color intensity (OD_595_) for 1 hour, and the relative azo dye reduction rate was calculated by setting the initial azo dye reduction rate in Δ*ispA* as 100%. E) Cu^2+^ reduction-induced toxicity reflected by growth inhibition. Growth of strains in GM17 media was monitored in a Bioscreen at 30 °C in absence and presence of 300 nM CuCl_2_, with initial OD = 0.1. The time to reach (TTR) OD = 0.4 was determined for each strain and condition, and the factor difference between the TTR in presence of 300 nM CuCl_2_ and in absence of CuCl_2_ is an indicator of growth inhibition, which reflects Cu^2+^ reduction. A) – D) data from three independent experiments, and E) data from biological triplicates. *shows significant (p<0.05) difference to MG1363. Error bars show SEM. AN: anaerobic; AE: aerobic; RES: respiration-permissive, i.e. heme and aeration.

The most distinct phenotypes among the mutants under the anaerobic conditions were observed when subjecting the mutants to an azo dye reduction (decolorization) and a Cu^2+^ reduction test. In these tests, indicators for the functionality of the different forms of MKs in *L. cremoris* EET, which are mostly described for anaerobic conditions, were obtained. In the azo dye reduction test, *ΔispA* showed the highest rate of decolorization among all strains (Fig. 3D, and example pictures of azo dye discoloration are provided in supplementary Fig. S4). The reduction rates for strains MG1363, *Δllmg_0196* and *ΔmenF* were 65%, 75% and 30% as compared to *ΔispA*. Observations in line with the azo dye decolorization test were obtained from the Cu^2+^ reduction test, which was reflected by the degree of growth inhibition in presence of Cu^2+^ (Fig. 3E). Strain *ΔmenF* was least inhibited in growth (factor 2) and *ΔispA* was most inhibited (factor 8), while wildtype strain MG1363 and the MK-3 producer *Δllmg_0196* showed an intermediate degree of growth inhibition (factor 4) under anaerobic conditions.

The primary metabolite profiles of MG1363 and mutants were mostly similar among each other under anaerobic conditions: all strains produced mainly lactate (ca. 55 mM) and hardly any acetate or acetoin (Fig. 4A). The analysis of the succinate concentration in culture supernatants revealed that *ΔispA* produced noticeable amount of succinate (0.5 mM) while in cultures of the other strains succinate was not detectable indicating consumption of the small amount of succinate present in M17 media, and this effect is most pronounced under anaerobic conditions (Fig. 4B).

**Figure 4.**
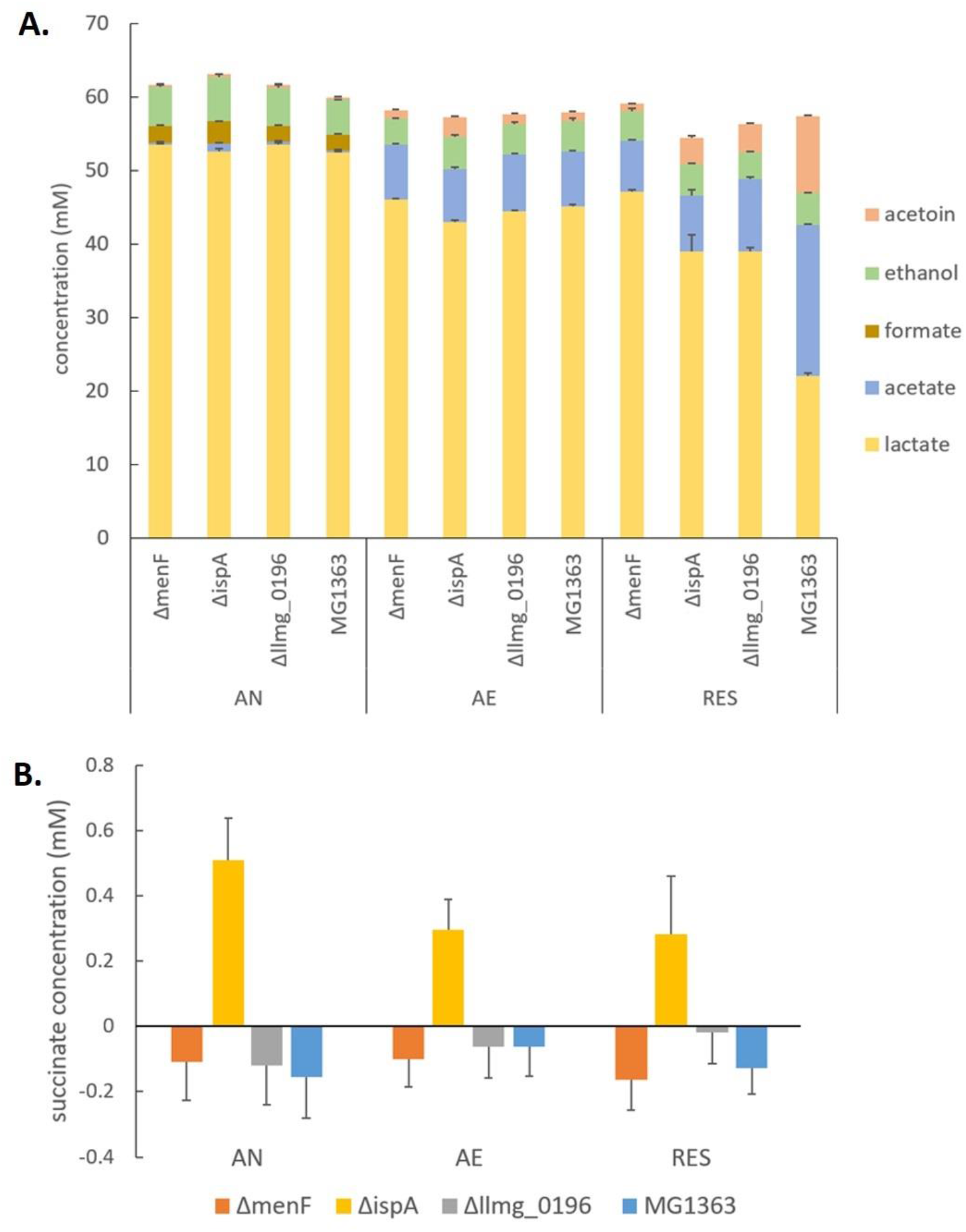
Metabolites in MG1363 and mutants. A) Concentration of primary metabolites including lactate, acetate, formate, acetoin and ethanol in the culture supernatant. B) Concentration of succinate in the culture supernatant. Metabolites were from the supernatant of cell cultures incubated in GM17 at 30 °C for 16-18 hours under indicated conditions. The concentration of metabolites in uncultured GM17 was used as level zero. A minor amount of succinate is present in GM17, and negative values from samples indicate consumption of the succinate present in GM17. AN: anaerobic; AE: aerobic; RES: respiration-permissive, i.e. heme and aeration. Data from three independent experiments, error bars show SEM.

### Aerobic conditions: mutants showed distinct phenotypes in long-term survival and Cu^2+^ reduction

Under aerobic conditions, all mutants showed similar biomass accumulation as compared to the wildtype strain MG1363 after overnight cultivation. The biomass yields were found to be at the same level as those observed under anaerobic conditions (Fig. 3A, supplementary Fig. S2A). Also the oxygen consumption rates for all strains were similar, ranging between 80 and 100 nmol/(min x OD unit) (Fig. 3B). The primary metabolite profiles of all strains cultivated under aerobic conditions were also similar to each other, with a small decrease in lactate (by 5 mM) and an increase in acetate (by 5 mM) concentration as compared to the anaerobic conditions (Fig. 4A).

The tested strains showed distinct phenotypes in survival when cultivated for a prolonged period of time under aerobic conditions (Fig. 3C): the presumed MK-1 producer *ΔispA* showed 1 log lower viability than the other strains after 24 h cultivation, and the difference even further increased after 48 h. While the culture of the non-MK producer *ΔmenF* remained at a viable plate count of 10 log CFU/mL, for strain *ΔispA* this value was lowered to 4 log CFU/mL, followed by the MK-3 producer *Δllmg_0196* and MK-9, 8, 3 producer MG1363 (6-7 log CFU/mL). After 72 h, none of the strains showed detectable viability (below 2 log CFU/mL) except strain *ΔmenF* which stood out with 8 log CFU/mL.

Big differences among the strains were also observed when the reduction of Cu^2+^ was tested under aerobic conditions (Fig. 3E). The growth inhibition, as reflected by the factor difference in the time it took the bacterial culture to grow from OD_600_ of 0.1 to 0.4 in absence and presence of Cu^2+^, was used to indicate Cu^2+^ reduction as the reduced product Cu^+^ poses toxicity to the cells. The Cu^2+^ reduction induced growth inhibition was most severe with strain *ΔispA* (factor 11), even more than under anaerobic conditions. While *ΔmenF* remains the least inhibited (factor 2), *Δllmg_0196* showed less inhibition (factor 3) than under anaerobic conditions, followed by MG1363 (factor 4).

### Respiration-permissive conditions: mutants showed distinct phenotypes in biomass accumulation, oxygen consumption and metabolite profiles

Under respiration-permissive conditions, differences were observed among the tested strains in biomass accumulation after overnight cultivation (Fig. 3A, supplementary Fig. S2A). The MK-9, 8, 3 producer MG1363 reached about 1.7 g/L in cell dry weight, which is almost a doubling compared to the yield found in anaerobic and aerobic conditions. In the non-MK producer Δ*menF* and presumed MK-1 producer *ΔispA*, the biomass accumulation remained the same level (1 g/L dry weight) as in anaerobic and aerobic conditions. In the MK-3 producer *Δllmg_0196*, about 1.3 g/L dry weight was obtained under respiration-permissive conditions, which is significantly higher than in Δ*menF* and *ΔispA* but lower than MG1363. When the mutants were complemented with the respective genes, the biomass accumulations became similar to strain MG1363 (supplementary Fig. S3). The stationary phase survival for all strains under the respiration-permissive conditions were not significantly different throughout 72 h, all maintained at about 9 log CFU/mL (now shown).

The oxygen consumption rates in strain MG1363 and mutants under respiration-permissive conditions also showed differences. While *ΔmenF* and *ΔispA* remained at the same level as in aerobic conditions (80-100 nmol/(min x OD unit)), MG1363 and *Δllmg_0196* showed much higher oxygen consumption rates, reaching values over 200 nmol/(min x OD unit).

Less difference among the strains were observed for the Cu^2+^ reduction test under the respiration-permissive conditions, all strains showed no growth inhibition caused by Cu^2+^ reduction (factor 1) except for *ΔispA* which still showed an inhibition factor of 5.

The metabolite profiles of strain MG1363 and mutants differed the most under respiration-permissive conditions (Fig. 4A). Strain MG1363 produced the lowest concentration of lactate (ca. 20 mM) and the highest acetate (20 mM) and acetoin (10 mM) concentrations among all strains. The metabolite profiles of *ΔmenF* were similar under the respiration-permissive and aerobic conditions. For *ΔispA* and *Δllmg_0196*, about 40 mM lactate and ca. 10 mM acetate, 5 mM acetoin was observed under respiration-permissive conditions, showing differences in comparison to their own profiles under aerobic conditions. Moreover, the profiles of strains *ΔispA* and *Δllmg_0196* also differed largely from that of strain MG1363 under respiration-permissive conditions.

### Additional insights provided by proteomics analysis

Finally, we examined the proteome profiles of strain MG1363 and mutants under anaerobic, aerobic and respiration-permissive conditions to further understand the phenotypic differences, which likely result from differences in bacterial metabolism, physiology and functionality caused by the short-chain and long-chain MK forms. Cells from stationary phase (16 h) were used for this analysis, and 1394 proteins in the sample set were quantified.

We examined changes in the proteome profile between the wildtype strain MG1363 and the MK mutants under the various cultivation conditions (Fig. 5). The proteome profiles obtained under anaerobic and aerobic conditions of cultures of the non-MK producer *ΔmenF* and the MK-3 producer *Δllmg_0196* were found to be very similar to that of strain MG1363. Under respiration-permissive conditions, *Δllmg_0196* showed a proteome profile similar to that of strain MG1363. In the comparison with strain MG1363, the presumed MK-1 producer *ΔispA* showed the most different proteome profile in all three conditions.

**Figure 5.**
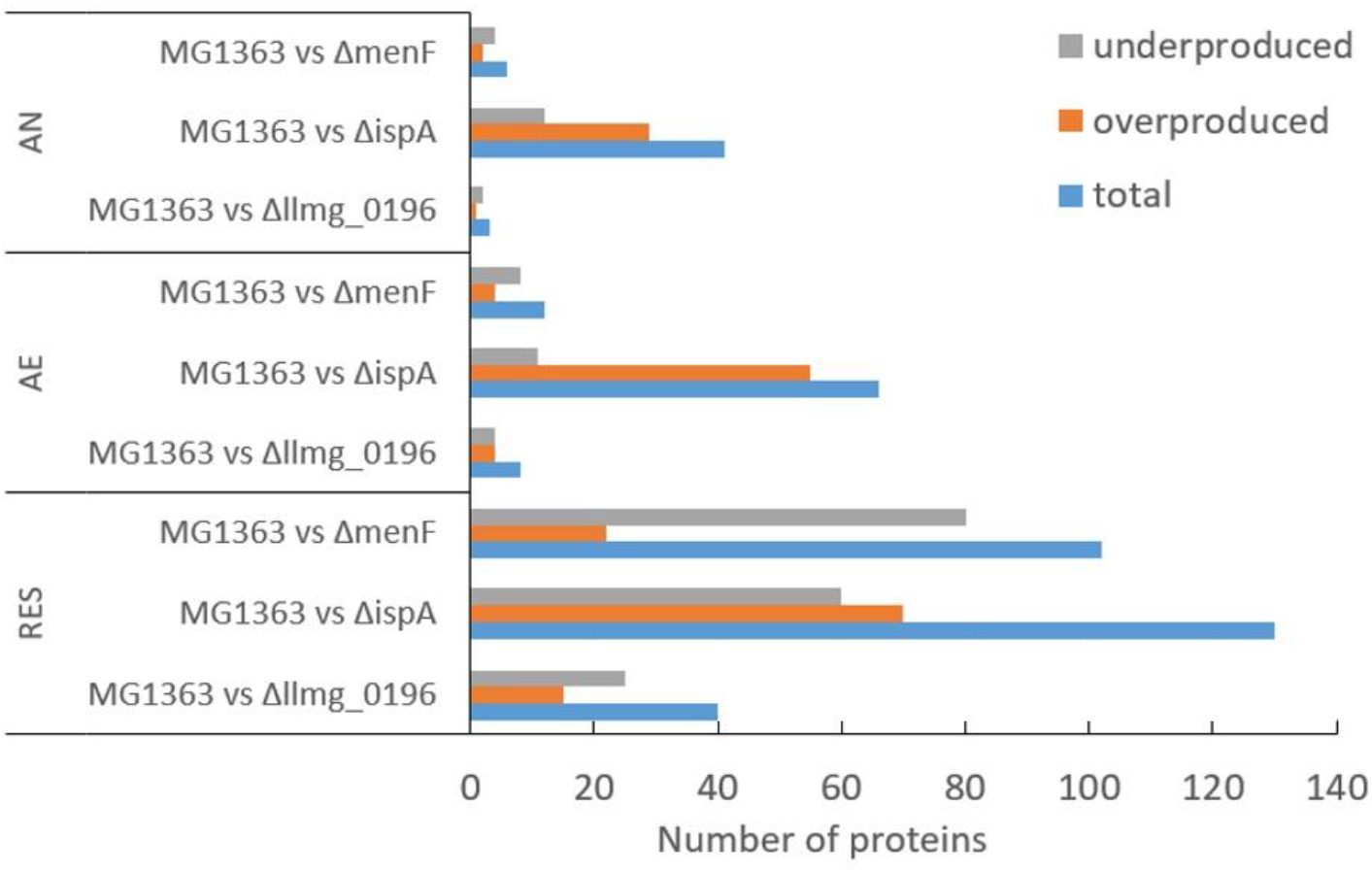
Differential proteome. Numbers of differentially produced proteins between MG1363 and mutants under each cultivation condition. Cells were obtained from cultures incubated in GM17 media at 30 °C for 16-18 hours under indicated conditions, samples from three independent experiments. AN: anaerobic; AE: aerobic; RES: respiration-permissive, i.e. heme and aeration. Proteins considered to be differently produced were selected when the LFQ intensity showed fold change ≥ 2 and p ≤ 0.05.

When closely examining the protein list, we identified proteins that are predicted to be involved in MK biosynthesis, aerobic respiration, aerobic growth and EET/anaerobic respiration (Table 1 – 4). Among proteins predicted to be part of the MK biosynthesis pathway (Table 1), we could first confirm that in respective mutants, the protein products of the deleted genes were indeed absent/only showed signals at very low levels possibly due to cross contaminations in sample preparations and quantification errors. Most other MK biosynthesis proteins in this list showed constitutive presence in all strains and conditions, and for the protein product (A2RM75) of *menB* for example, a slightly higher level was observed under the aerobic conditions, which could contribute to the higher MK content in *L. cremoris* under aerobic conditions observed in this study (Fig. 2). The protein product (A2RHR6) of the gene annotated as *menA* was not detected in all cases, but the homologous protein (A2RJB1) encoded by *ubiA* was present constitutively, suggesting that the gene annotated as *ubiA* is in fact functional in merging the head and tail group of MKs in strain MG1363 instead of the gene annotated as *menA*. The absence of protein product of *gerCA* (gene deletion mutant of *gerCA*) and *ispB* showed no influence on the MK profile) and inconsistent detection of *menX* product, also highlighted the necessity to experimentally confirm the functionality of the genes predicted in the MK biosynthesis pathway.

**Table 1.**
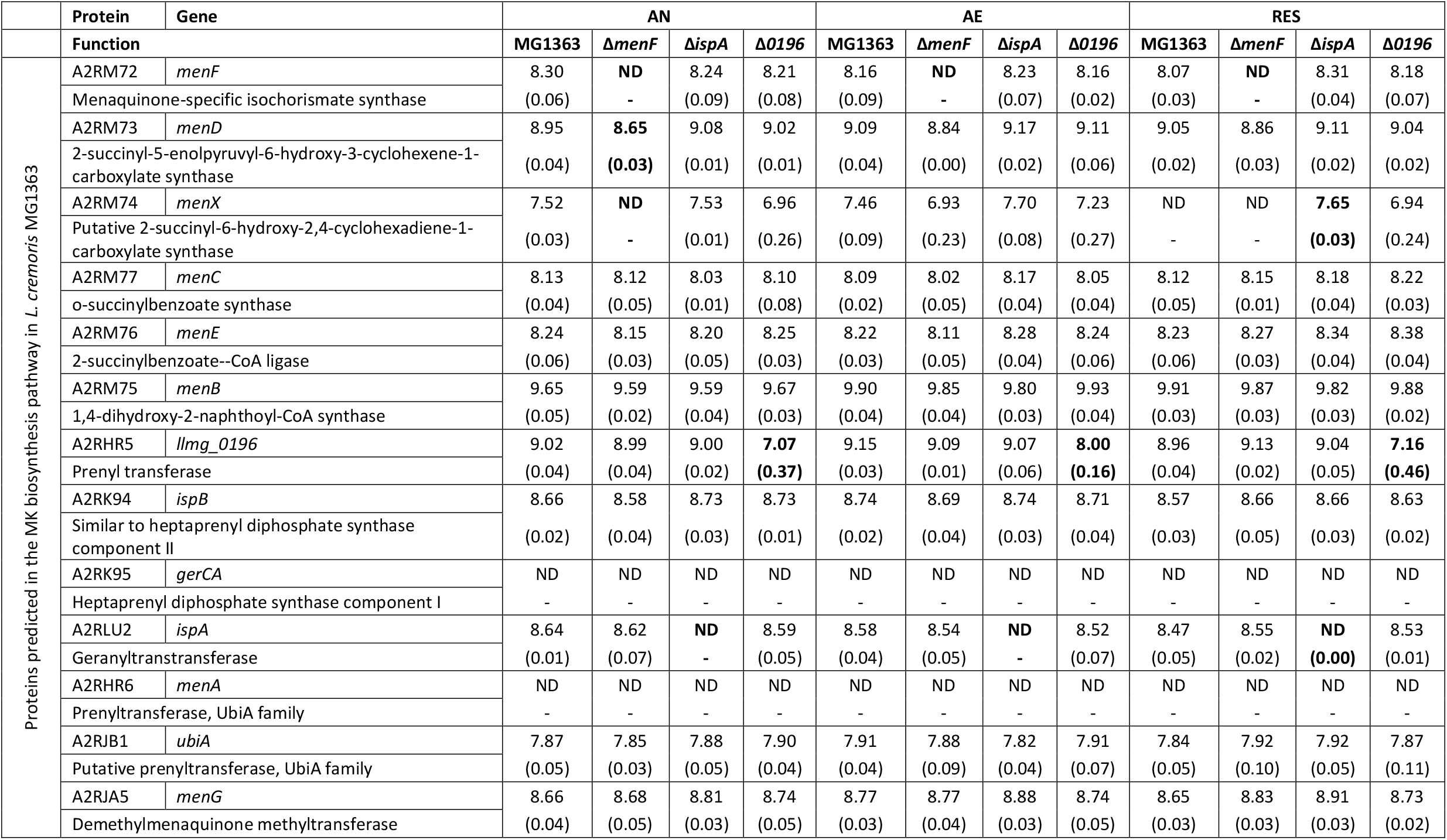
Quantity (Log LFQ intensity) of proteins predicted in the MK biosynthesis pathway in MG1363 and mutant under different cultiv ation conditions. Values are average from samples collected from 3 independent experiments, SEM values are shown in brackets. Detection limit in Log LFQ intensity: 6.7; Δ*0196* = Δ*llmg_0196*. Values in mutants that are significantly different (p < 0.05, fold change > 2) from strain MG1363 under the same cultivation conditions are highlighted in bold letters.

**Table 2.** Quantity (Log LFQ intensity) of proteins involved in aerobic respiratory ETC in MG1363 and mutant under different cultivation conditions. Values are average from samples collected from 3 independent experiments, SEM values are shown in brackets. Detection limit in Log LFQ intensity: 6.7; Δ*0196* = Δ*llmg_0196*. Values in mutants <u>that are significantly different (p < 0.05, fold change > 2) from strain MG1363 under the same cultivation conditions are highlighted in bold letters.</u>

|  | Protein | Gene | AN |  |  |  | AE |  |  |  | RES |  |  |  |
| --- | --- | --- | --- | --- | --- | --- | --- | --- | --- | --- | --- | --- | --- | --- |
| | Function | | MG1363 | $\Delta menF$ | $\Delta ispA$ | $\Delta 0196$ | MG1363 | $\Delta menF$ | $\Delta ispA$ | $\Delta 0196$ | MG1363 | $\Delta menF$ | $\Delta ispA$ | $\Delta 0196$ |
| Proteins involved in aerobic respiratory ETC | A2RLY0 | *noxB* | 10.02 | 10.00 | 10.01 | 10.02 | 10.01 | 9.97 | 10.05 | 10.03 | 9.93 | 10.00 | 10.04 | 9.97 |
|  | NADH dehydrogenase |  | (0.04) | (0.02) | (0.03) | (0.01) | (0.03) | (0.02) | (0.03) | (0.03) | (0.01) | (0.01) | (0.02) | (0.05) |
|  | A2RLY1 | *noxA* | 9.70 | 9.54 | 9.50 | 9.56 | 9.73 | 9.61 | 9.61 | 9.65 | 9.59 | 9.61 | 9.64 | 9.62 |
|  | NADH dehydrogenase |  | (0.03) | (0.03) | (0.04) | (0.05) | (0.04) | (0.02) | (0.02) | (0.02) | (0.03) | (0.02) | (0.02) | (0.04) |
|  | A2RHR4 | *llmg_0195* | 9.65 | 9.63 | 9.62 | 9.51 | 9.58 | 9.57 | 9.56 | 9.49 | 9.63 | 9.54 | 9.54 | 9.60 |
|  | Putative NADH dehydrogenase |  | (0.03) | (0.02) | (0.03) | (0.08) | (0.03) | (0.02) | (0.04) | (0.03) | (0.02) | (0.02) | (0.04) | (0.04) |
|  | A2RMA6 | *cydB* | 8.76 | 8.77 | 8.65 | 8.69 | 8.65 | 8.58 | 8.51 | 8.62 | 8.47 | 8.60 | 8.45 | 8.69 |
|  | Cytochrome d ubiquinol oxidase, subunit II |  | (0.06) | (0.11) | (0.08) | (0.12) | (0.02) | (0.05) | (0.11) | (0.05) | (0.05) | (0.11) | (0.04) | (0.10) |
|  | A2RMA7 | *cydA* | 8.71 | 8.85 | 8.75 | 8.86 | 8.85 | 8.73 | 8.62 | 8.84 | 8.72 | 8.79 | 8.52 | 8.72 |
|  | Cytochrome bd-I ubiquinol oxidase subunit I |  | (0.14) | (0.09) | (0.05) | (0.02) | (0.09) | (0.01) | (0.08) | (0.02) | (0.07) | (0.01) | (0.03) | (0.06) |
|  | A2RIR7 | *hemK* | 8.22 | 8.16 | 8.27 | 8.15 | 7.99 | 7.83 | 8.24 | 7.84 | 7.81 | 7.95 | **8.29** | 7.84 |
|  | Release factor glutamine methyltransferase |  | (0.03) | (0.03) | (0.01) | (0.01) | (0.05) | (0.05) | (0.03) | (0.04) | (0.02) | (0.04) | **(0.05)** | (0.03) |
|  | A2RL35 | *hemN* | 7.67 | 7.67 | 7.67 | 7.68 | ND | **7.44** | **7.50** | **7.44** | 7.46 | 6.93 | 7.59 | 7.13 |
|  | Coproporphyrinogen III oxidase |  | (0.05) | (0.06) | (0.07) | (0.04) | - | **(0.14)** | **(0.03)** | **(0.06)** | (0.03) | (0.23) | (0.03) | (0.22) |

**Table 3.**
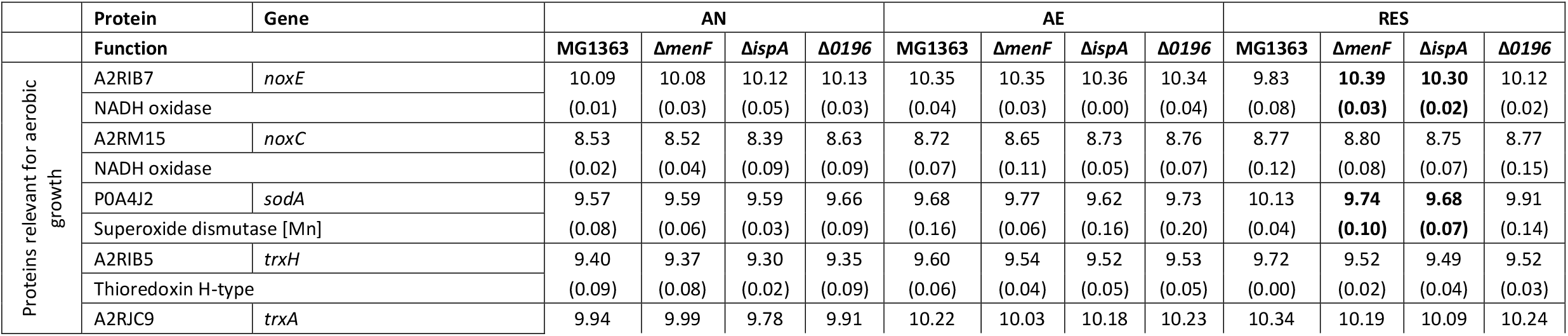
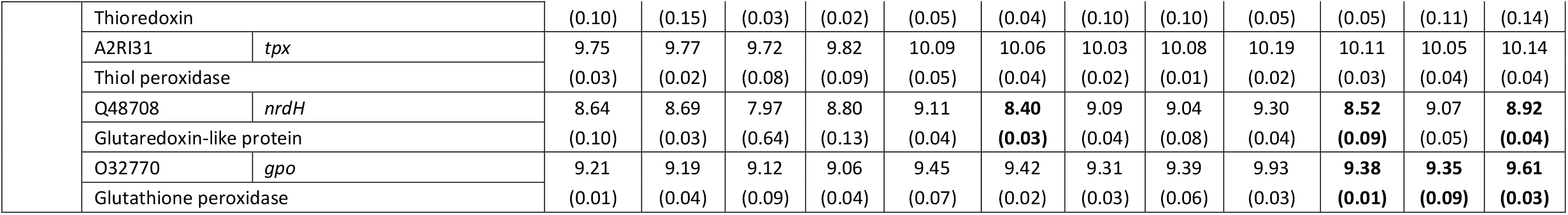
Quantity (Log LFQ intensity) of proteins relevant for aerobic growth in general in MG1363 and mutant under different cultivation conditions. Values are average from samples collected from 3 independent experiments, SEM values are shown in brackets. Detection limit in Log LFQ intensity: 6.7; Δ*0196* = Δ*llmg_0196*. Values in mutants that are significantly different (p < 0.05, fold change > 2) from strain MG1363 under the same cultivation conditions are highlighted in bold letters.

**Table 4.**
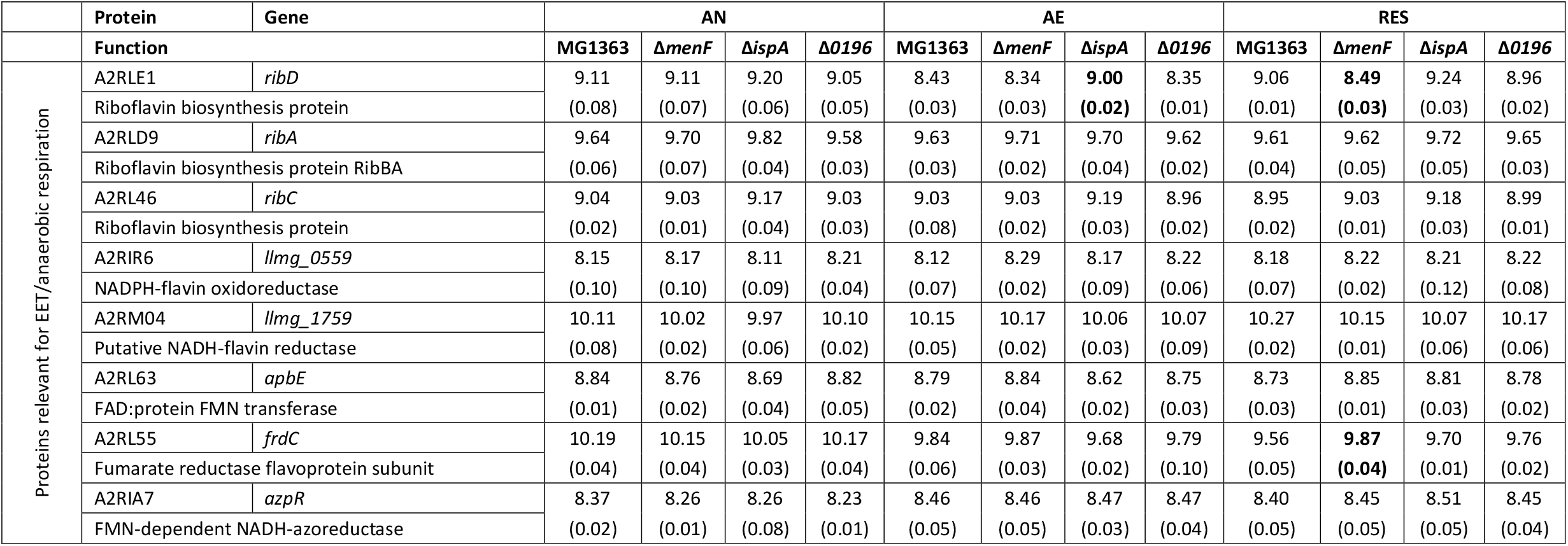
Quantity (Log LFQ intensity) of proteins relevant for EET/anaerobic respiration in MG1363 and mutant under different cultivation conditions. Values are average from samples collected from 3 independent experiments, SEM values are shown in brackets. Detection limit in Log LFQ intensity: 6.7; Δ*0196* = Δ*llmg_0196*. Values in mutants that are significantly different (p < 0.05, fold change > 2) from strain MG1363 under the same cultivation conditions are highlighted in bold letters.

When examining the proteins predicted to be involved in aerobic respiration (Table 2), including the NADH dehydrogenase and cytochrome *bd* oxidase in the respiratory ETC, we see that these proteins are present in all strains under all cultivation conditions at similar levels. Protein products of two heme traffic/synthesis genes *hemN* and *hemK* were constantly present in most cases. Two type II NADH dehydrogenases have been identified in the proteome, encoded by genes *noxA* and *noxB*.

We also examined other proteins relevant for the aerobic cultivation conditions (Table 3), including NADH oxidase and several oxidoreductases. In general proteins in this category were produced at higher levels under the aerobic conditions than anaerobic, as expected. The NADH oxidase (A2RIB7) in MG1363 and Δ*llmg_0196* was produced at a lower level in respiration-permissive conditions compared to aerobic conditions, which could be a result of the activated respiratory ETC taking over the reduction of oxygen in these two strains. Antioxidation proteins like thioredoxins (A2RIB5, A2RJC9, A2RI31) were in general more abundant under the aerobic conditions than anaerobic conditions. Moreover, the low level of glutaredoxin-like protein (Q48708) in the Δ*menF* under the aerobic conditions indeed reflects particular roles of MKs under oxidative/aerobic conditions.

Furthermore, we examined proteins predicted to be relevant for EET/anaerobic respiration (Table 4). While the NADH dehydrogenase encoded by *noxB* and most flavin synthesis or oxidoreduction related proteins were constantly present at similar levels in all strains and conditions, one riboflavin biosynthesis protein (A2RLE1) was more abundant in the MK-1 producer Δ*ispA* than other strains under aerobic and respiration-permissive conditions. A fumarate reductase flavoprotein subunit protein (A2RL55) showed the highest level under anaerobic conditions in all strains, pointing to a potential role in anaerobic respiration.

Besides the proteins that are expected to be highly relevant for the several cultivation conditions and electron transfer pathways, much more information could be derived from the proteomics analysis. For example, when examining the proteins involved in primary metabolism (supplementary Table S4), we observed protein level changes across the three cultivation conditions in the tested strains, which corresponds to the production of primary metabolites (Fig. 4A). When examining the remainder of the proteins that showed significant changes across the strains or cultivation conditions, membrane proteins, oxidoreductases, stress proteins, metal ion (copper, iron, etc.) transporters, ATP-binding cassette (ABC) transporters were among the most often observed categories. A non-exhaustive list of these proteins is provided in supplementary Table S5.

## Discussion

*L. cremoris* and *L. lactis* are best-known for their occurrence and applications in dairy fermentations, but their niche extends to a range of natural and food production environments, where the bacteria encounters different growth conditions including anaerobic, aerobic and respiration-permissive conditions. *L. cremoris* and *L. lactis* are among the few LAB species that maintained the ability to produce MKs (vitamin K2), in both short-chain (MK-3) and long-chain forms (MK-9, MK-8). The physiological significance of the short-chain and long-chain MK forms in the lifestyle of *L. cremoris* and *L. lactis* has not been investigated extensively. In this study, we constructed *L. cremoris* mutants producing short-chain and long-chain MKs: deletion of genes *menF, ispA* and *llmg_0196* resulted in a non-MK producer, a presumed MK-1 producer and a MK-3 producer, respectively. Together with the wildtype MG1363 producing mainly MK-9 and MK-8, these mutants with gene deletions served to further elucidate the functionality of MKs in *L. cremoris*. By exposing this set of strains to anaerobic, aerobic and respiration-permissive conditions, functionalities of long-chain and short-chain MKs in *L. cremoris* under different environmental conditions could be inferred (Fig. 6).

**Figure 6.**
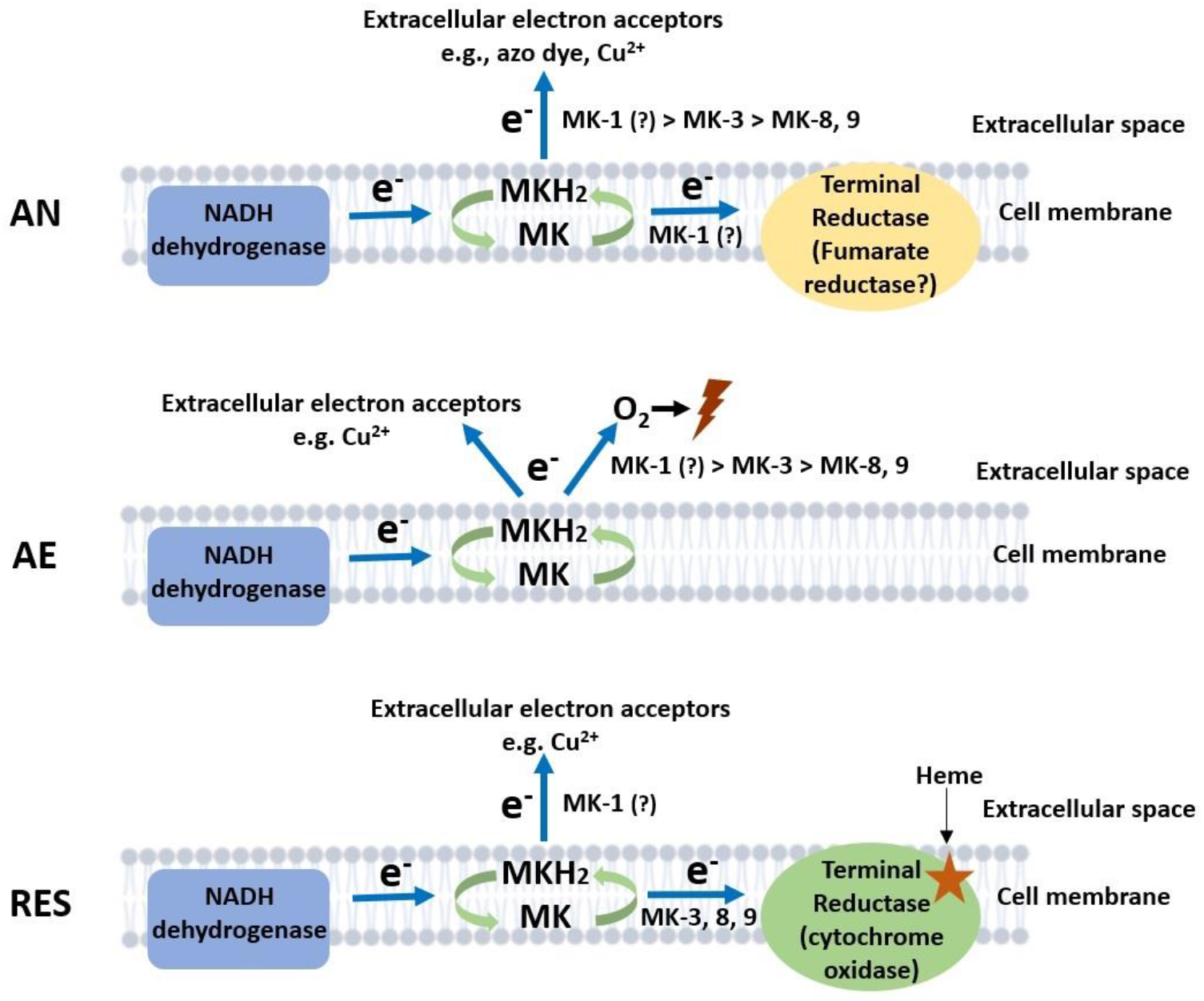
Proposed model of electron flows facilitated by MKs under anaerobic, aerobic and respiration-permissive conditions. The predominantly involved MK forms are annotated next to the arrows indicating electron flows, with “>” symbols indicate preferences. Lightning symbol represents loss of viability in the cells. AN: anaerobic; AE: aerobic; RES: respiration-permissive, i.e. heme and aeration. Note that the illustration is only a schematic presentation of proposed electron flow routes, the location of enzymes are not confirmed in this study.

We could confirm that *llmg_0196* indeed encodes a (nona)prenyl diphosphate synthase as the corresponding gene deletion mutant only produced MK-3. The level of MK-3 in Δ*llmg_0196* was lower than the total MK level in MG1363, indicating that there are feedback mechanisms influenced by the intermediate compounds of the MK biosynthetic pathway. Another possible explanation to this is a lower efficiency of the DHNA polyprenyltransferase (MenA) in incorporating the farnesyl diphosphate group (containing 3 isoprene units) to the head group.

Based on the observed MK profiles alone, it can be concluded that strains *ΔmenF* and *ΔispA* hardly produce MKs. The remaining low amount of MK-9 produced in strain *ΔispA* could be a result of additional activity of the prenyl diphosphate synthase encoded by *llmg_0196*. Combining other analyses that demonstrated the role of MKs, we saw distinct phenotypes of *ΔmenF* and *ΔispA*: in *ΔmenF*, phenotypes from all processes including aerobic respiration or copper reduction suggested absence of MK-related functions. In *ΔispA*, clear activities of e.g. azo dye/copper reduction could be observed, suggesting presence of (a) MK form(s) that was not detected in the MK profile analysis. It is most plausible that MK-1 was produced by *ΔispA* but was not revealed in the analysis, because the MK extraction method used in this study is based on hexane extraction of hydrophobic/lipophilic molecules. This method works well for long-chain MKs and lipids, but it is likely that MK-1 is too low in hydrophobicity to be extracted by hexane from the aqueous environment. Notably, this extraction method is commonly adopted for MK analysis in scientific research, which could mean that the presence and role of MK-1, if any, may have been missed so far in MK related studies.

Nevertheless, we could infer the function of MK-1 in the current study from the different phenotypes of the mutants. In this context it is worth mentioning that, we did not analyze the presence of any MK precursors such as demethylmenaquinones DMKs, previously studied in *L. cremoris* (Rezaïki et al., 2008) and *Listeria monocytogenes* (Light et al., 2018). Here we focused on the MK forms with various side-chain length, and will therefore not discuss the effect of the methylation (DMK to MK) separately. Moreover, deletion of gene *ispA* as reported for other bacteria is expected to display pleiotropic effects including impaired growth (Krute et al., 2015). This was also observed for strain *ΔispA* in this study since *ΔispA* shows a slightly lower growth rate in comparison to the other strains (supplementary Fig. S2B).

Based on existing knowledge on MK roles in bacteria (Duwat et al., 2001; Koebmann et al., 2008; Rezaïki et al., 2008; Light et al., 2018), we examined the phenotypes of mutants in anaerobic, aerobic and respiration-permissive conditions. The common or different functional features of short-chain MKs and long-chain MKs were revealed in the three conditions (Fig. 6).

Under anaerobic conditions, the most distinct phenotypes among the strains were the ability to reduce azo dye and copper ions, which are indications of EET. The EET efficiency appeared to be higher in the presumed MK-1 producer *ΔispA* than the MK-3 and long-chain MKs producers, indicating a preference in anaerobic EET for short-chain MKs over long-chain MKs (Fig. 6). The same preference was suggested by the metabolite analysis where succinate was formed by *ΔispA*, plausibly as a result of anaerobic electron transport chain with fumarate serving as the electron acceptor. The role of MKs in anaerobic respiration via reduction of fumarate to succinate has been reported for *E. coli* and *En. faecalis* (Huycke et al., 2001; Brooijmans et al., 2009a). Genome annotation of *L. cremoris* strain MG1363 indeed reveals the gene *frdC* encoding a fumarate reductase, homologous to the fumarate reductases reported for *E. coli* and *En. faecalis*. In fact, activity of a fumarate reductase was indeed reported in *L. lactis* a long time ago (Hillier et al., 1979). In agreement with the theory, a fumarate reductase flavoprotein subunit protein (A2RL55) was also identified in this study in the proteome of all strains, with high abundance under anaerobic conditions. In addition, the higher stationary phase survival of *ΔispA* and Δ*llmg_0196* under anaerobic conditions supports the hypothesis that short-chain MKs offer an advantage for *L. cremoris* in anaerobic respiration/EET, allowing additional energy generation for maintenance in the stationary phase cells.

Under aerobic conditions, large differences were observed in stationary phase survival of the strains, where non-MK producer *ΔmenF* maintained high viability throughout the incubation period while the presumed MK-1 producer *ΔispA* lost viability fastest followed by the MK-3 producer Δ*llmg_0196*. Under aerobic conditions where the cytochrome oxidase is not functional (no heme supplementation), MKs have been suggested to mediate the reduction of oxygen, promoting ROS formation in *L. cremoris* (Rezaïki et al., 2008). The MK-mediated ROS formation could reduce the viability of cells, explaining the observed differences in the tested strains. In the proposed model, we assume MK-mediated ROS formation as an extracellular activity. This model is supported by measurement of extracellular superoxide formation mediated by (D)MKs in *L. cremoris* (Rezaïki et al., 2008) and *En. faecalis* (Huycke et al., 2001) under aerobic conditions. However, we do not exclude the possibility that MK-mediated ROS formation also takes place in the cytoplasm, but this intracellular ROS fraction is not expected to pose major damage to the cells due to the activity of an intracellular superoxide dismutase (Huycke et al., 2001; Ballal et al., 2015), which has also been identified in the proteome determined in our study (Table 3, protein ID P0A4J2). Therefore, the difference in cell viability observed after prolonged cultivation under aerobic conditions is considered to be mainly a result of the extracellular ROS formation mediated by MKs. Here a preference for short-chain MKs seemed to exist for the reaction with oxygen/ROS formation too (Fig. 6).

Furthermore, the copper reduction still took place under aerobic conditions, but to a lesser extent in *Δllmg_0196* as compared to anaerobic conditions. This could mean that, in absence of a functional cytochrome oxidase, the electron flow via MKs towards oxygen and EET can both be active, with possible competition between the two pathways of the electrons (Fig. 6).

Under respiration-permissive conditions, it was consistently shown that MG1363 demonstrated all aerobic respiration-related phenotypes: increased biomass yield in comparison to fermentative growth due to more efficient ATP generation, higher oxygen consumption rate than under aerobic conditions, large decrease in lactate production and increase in acetate and acetoin as compared to cultures in the fermentative growth mode as a result of redox cofactor (NAD/NADH) changes caused by the membrane embedded ETC, etc. (Duwat et al., 2001; Brooijmans et al., 2009c). The MK-3 producer *Δllmg_0196* demonstrated these phenotypical changes, but to a lesser extent than MG1363, which could be explained by the lower content of MKs. The non-MK producer *ΔmenF* as well as presumed MK-1 producer *ΔispA* did not shown these changes. The proteome profiles reflected the same change/difference among the strains. It could be collectively deduced that long-chain MKs (MK-9 and MK-8) and MK-3, but not MK-1, complement the respiratory ETC in *L. cremoris*.

As a functional cytochrome oxidase was provided, the copper reduction activity was attenuated in MG1363 and *Δllmg_0196*. This observation agrees with the previous findings: heme-induced respiration of *L. lactis* suppressed copper reduction (Abicht et al., 2013); *L. cremoris* and *L. lactis* could decolorize azo dyes only under anaerobic conditions (Pérez-Díaz and McFeeters, 2009); the EET in *En. faecalis* was attenuated by cytochrome *bd* oxidase activity (Pankratova et al., 2018). The remaining copper reduction activity in *ΔispA* suggest that MK-1 has the highest affinity for EET among all studied electron transfer routes.

The different efficiency of long-chain and short-chain MKs in the various electron transfer pathways is considered to be a result of the hydrophobicity of the MK forms in relation to the location where electron transfer takes place. The respiratory ETC has all components located in the lipid bilayer of the cell membrane, and long-chain MKs are well-embedded in the membrane due to the strong hydrophobicity. In contrast, pathways like EET direct electrons towards the extracellular electron acceptors, and thus the short-chain MKs with higher hydrophilicity demonstrate advantages in transferring electrons to the free flavin shuttles (Light et al., 2018), or even act as soluble electron shuttles themselves (Freguia et al., 2009).

This study examined the functions of short-chain and long-chain MKs in *L. cremoris* in lab conditions, but the significance can be inferred for *L. cremoris* and *L. lactis* in natural settings too. For example, the role of MKs in EET allows more options for bacteria to utilize carbon sources that are otherwise not fermentable (Light et al., 2018). Although not investigated extensively in this study, we did identify a putative fumarate reductase flavoprotein (Table 4, protein ID A2RL55) that is overproduced under anaerobic conditions in *L. cremoris*. This fumarate reductase shows homology to a cell surface-associated fumarate reductase described in *Ls. monocytogenes* (Light et al., 2019). The fumarate reductase in *Ls. monocytogenes* was found to rely on EET to reduce fumarate to succinate, supporting growth of the bacterium via an anaerobic respiration mechanism; an EET mutant showed competitive disadvantage in the gut of mouse (Light et al., 2018). Similarly, competitive advantages offered by EET mediated by MKs can be inferred for *L. cremoris* and *L. lactis*: for the application in cheese production, the ripening process takes place when these bacteria face nutrient limitation/carbon starvation, but metabolic activity is still desired for flavor development in cheese (Smid and Kleerebezem, 2014). EET mediated by MKs, especially the short-chain MKs, are expected to offer advantage to *L. cremoris* and*L. lactis* for additional nutrient source utilization and maintaining longer survival/metabolic activity period, ideal for successful or even accelerated cheese ripening.

The possible role of MKs in ROS formation and the consequence for stationary phase survival of *L. cremoris* suggest disadvantages of MK production in this bacterium, in particular under aerobic conditions when heme is not supplied. This knowledge can be applied for selecting natural variants of *L. cremoris* and *L. lactis* that are extra resistant to the oxidative conditions during starter culture production and initial stages in cheese production, given the improved robustness of the non-MK producer under aerobic conditions. Interestingly, a previous study demonstrated that adaptive laboratory evolution of *L. cremoris* under highly aerated conditions resulted in evolved mutants showing elevated vitamin K2 content and enhanced resistance to oxidative stress (Liu et al., 2021). This supports that natural selection methods, based on the roles of MKs in *L. cremoris*, allow us to obtain different types of variants that are of interest for applications.

Nevertheless, as the effect of MKs in *L. cremoris* survival is only displayed after prolonged exposure to oxygen (48-72 h), MKs can be considered to be beneficial for *L. cremoris* in dairy fermentation, where the bacterium is more exposed to anaerobic conditions. In other natural settings, such as the original niche of *L. cremoris* and *L. lactis*, i.e., plant surfaces, heme and oxygen are often available (Pedersen et al., 2012; Cavanagh et al., 2015). Under such circumstances, maintaining the MK producing ability indeed offers advantage to *L. cremoris* and *L. lactis* as the aerobic respiration promotes the growth and survival of these bacteria (Duwat et al., 2001; Gaudu et al., 2002). The heme-induced respiration has been applied in starter production to obtain higher yield and more robust *L. cremoris* and *L. lactis* for dairy industry (Pedersen et al., 2005; Garrigues et al., 2006).

Notably, MKs are also known as vitamin K2, a fat-soluble vitamin essential to human health (Vermeer and Schurgers, 2000; Beulens et al., 2013). In the human body, vitamin K2 acts as the co-factor for γ-glutamyl carboxylase catalyzing the carboxylation of the glutamate residues in Gla-proteins, thereby ensuring biological functions of Gla-proteins in essential human physiological processes including hemostasis, calcium metabolism and cell growth regulation. Dietary intake of vitamin K2 has been associated with reduced risk of coronary heart disease and improved bone health (Geleijnse et al., 2004; Fujita et al., 2012). In terms of benefits for human health, the interest often goes to the long-chain MKs for the higher bioavailability they demonstrated compared to the short-chain MKs; the association with cardiovascular health was also demonstrated prominently with intake of long-chain MKs (Geleijnse et al., 2004; Sato et al., 2012). The insight in the physiological functions of MKs in *L. cremoris* also provides opportunities for vitamin K2 enrichment of fermented food products: suitable conditions can be designed accordingly for natural selection procedures to obtain variants with desired vitamin K2 production profiles.

Taken together, the set of both long-chain and short-chain MKs produced by *L. cremoris* (and *L. lactis*), in combination with the different functionality of MK variants in anaerobic, aerobic and respiration-permissive conditions, points to an efficient strategy to adapt to diverse aeration and nutritional conditions that *L. cremoris* and *L. lactis* strains may encounter in their natural niche or during food fermentation processes. It is highly desirable to extend the inferred significance of MKs in *L. cremoris* from lab settings to natural ecological systems with further investigations. As the short-chain MKs are more water soluble and allow possibilities for cross-feeding (Rezaïki et al., 2008), their relevance for interacting not only with the environment, but also with hosts or cohabitating microbes extends the significance to diverse habitats.

## Data availability statement

The datasets used and/or analyzed during the current study are available from the corresponding author on reasonable request.

## Authors’ contributions

YL, EJS and TA conceived the study. YL designed the experiments, executed experiments and carried out the data analysis and interpretation. NC contributed to the phenotypical characterization of the mutants. SB contributed to obtaining the proteomics data. NC, JB and AGL contributed to mutant construction. YL, EJS and TA wrote the manuscript.

All authors read and approved the final manuscript.

## Acknowledgements

The authors would like to thank Jeroen Koomen (Food Microbiology, Wageningen University) for his kind help in analyzing proteomics data. The authors also thank Mark Sanders (Food Chemistry, Wageningen University) for his assistance in vitamin K2 analysis.

