## Supplementary for "Physiological roles of short-chain and long-chain menaquinones (vitamin K2) in *Lactococcus cremoris*"

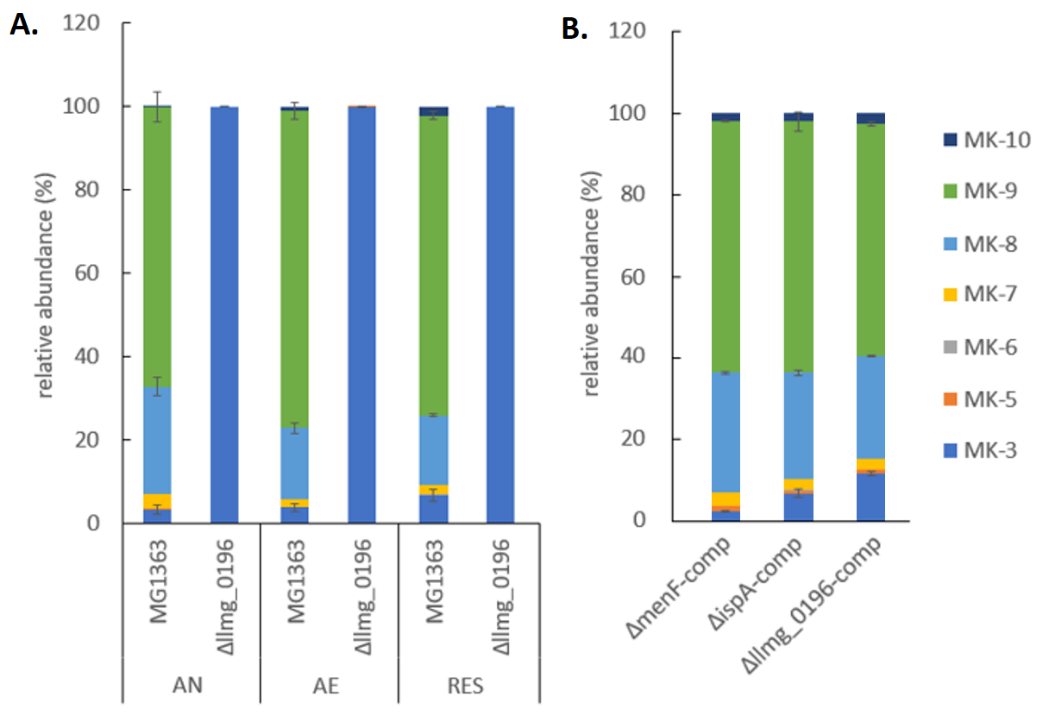

**Figure S1. MK profile in the biomass of MG1363 and mutants.** A) Relative abundance of tested MK forms in MG1363 and mutants with gene deletions. Note that only trace amount of MKs were detected in Δ*menF* and Δ*ispA*, and the relative abundance is therefore not shown. All strains were cultivated in GM17 medium under indicated conditions for 16-18 hours at 30 °C before MKs were extracted from the biomass. AN: anaerobic; AE: aerobic; RES: respiration-permissive, i.e. heme and aeration. Samples for each strain and condition were from three independent experiments. B) Relative abundance of tested MK forms in mutants with respective gene complementation. Strains were cultivated in GM17 medium under anaerobic conditions for 16-18 hours at 30 °C, 5 μg/mL erythromycin was added to the medium, data were from four biological replicates. Error bars show SEM for MK-3, MK-8 and MK-9.

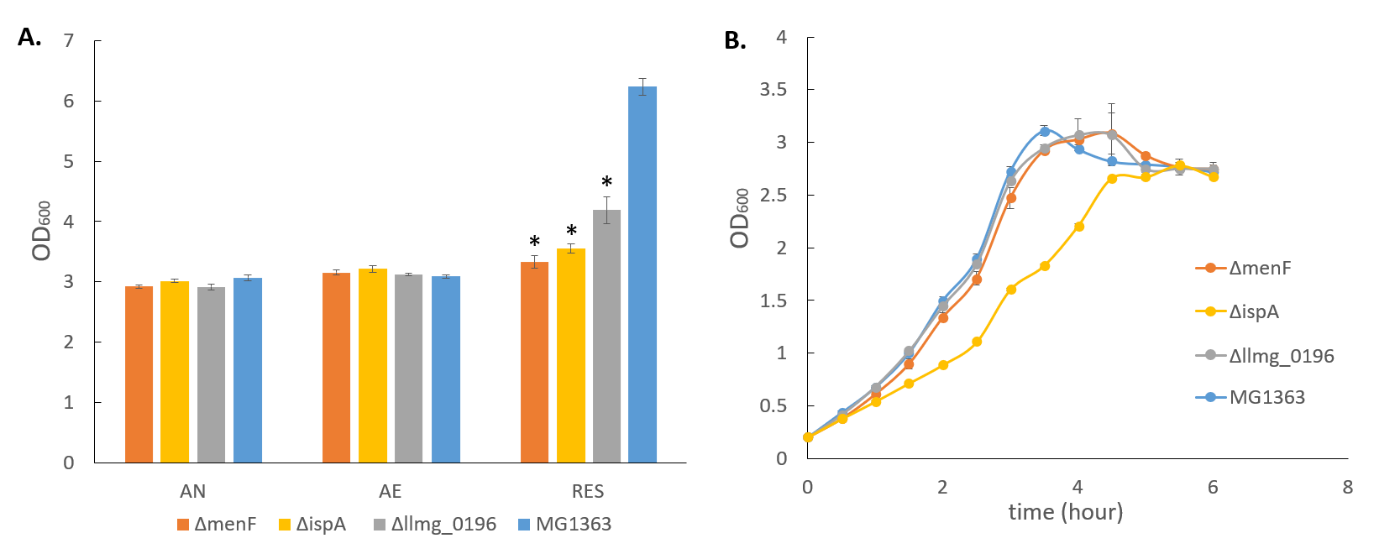

**Figure S2. Growth performance of MG1363 and mutants.** A) Biomass accumulation measured by OD. Data from four independent experiments. AN: anaerobic; AE: aerobic; RES: respiration-permissive, i.e. heme and aeration. *shows significant (p<0.05) difference to MG1363. B) growth curves. The growth was monitored under anaerobic conditions. Data from three independent experiments. Error bars show SEM.

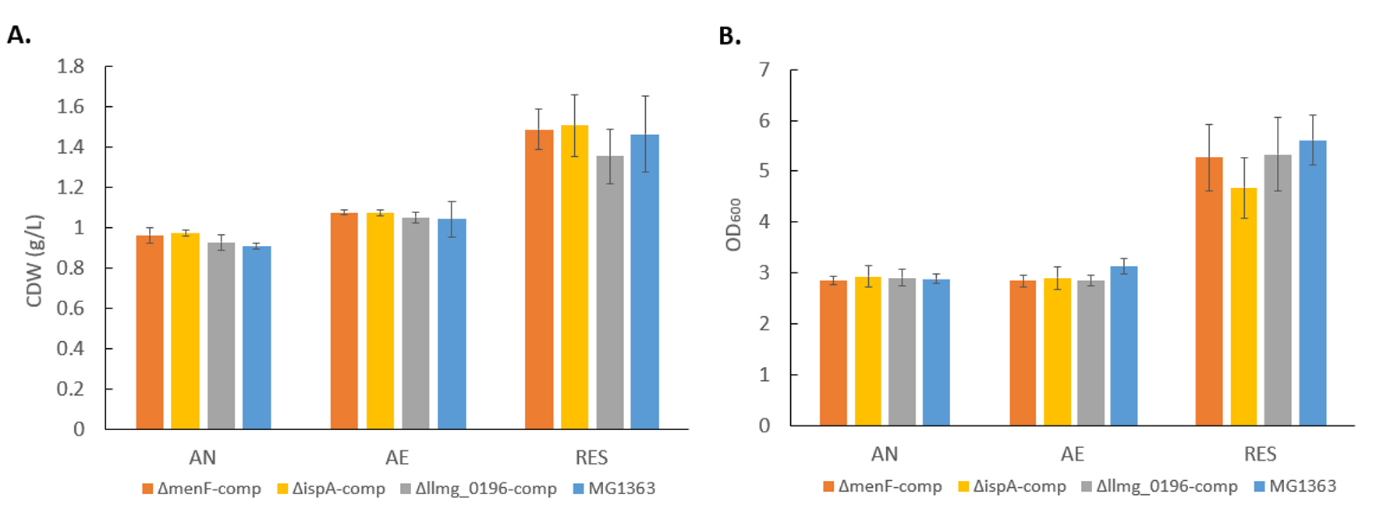

**Figure S3. Biomass accumulation of MG1363 and mutants with gene complementation.** A) Cell dry weight (CDW). Data from four independent experiments. B) OD measurement. Data from three independent experiments. Error bars show SEM. AN: anaerobic; AE: aerobic; RES: respiration-permissive, i.e. heme and aeration.

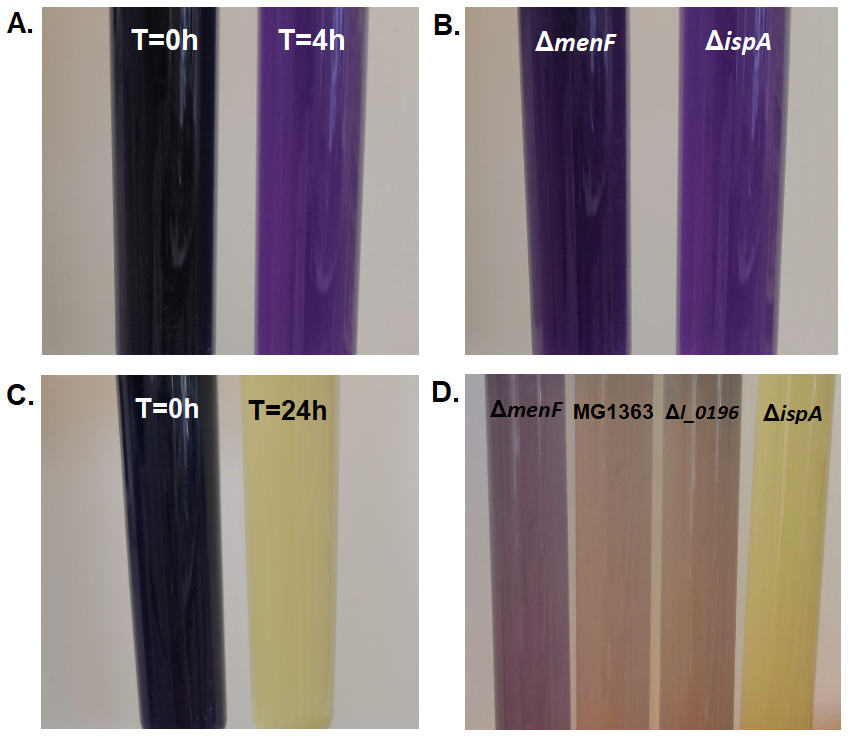

**Figure S4. Example pictures of azo dye decolorization by *L. cremoris*.** A) Reaction of *ΔispA* at 0 h and 4 h. B) Reaction of *ΔmenF* and *ΔispA* at 4 h. C) Reaction of *ΔispA* at 0 h and 24 h. D) Reaction of all strains at 24 h. Azo dye reduction was monitored in anaerobic conditions.

**Table S1. Primers used for PCR amplifying upstream and downstream homologous regions for the target genes.**

| Name^a^ | Primer sequence 5’ – 3’^b^ |
| --- | --- |
| 0196_HR1_HindIII_Fw | **AACGTGAAGCTTTACTGACAG** |
| 0196_HR1_PstI_Rv | CGCGCGCTGCAG**CAATGTTTTTCTCTTTCCTCTCG** |
| 0196_HR2_PstI_Fw | AAGAAGCTGCAG**TAAAAGCCTGTCACCGATAGG** |
| 0196_HR2_XbaI_Rv | GGCGGCTCTAGA**ATCACAGGTGTTATTCGCTAAC** |
| ispA_HR1_HindIII_Fw | CCACCAAAGCTT**CCCTCCTCGACTAACATATCG** |
| ispA_HR1_PstI_Rv | AAGAAGCTGCAG**GAACAACTGGAAGTGGAATAAATG** |
| ispA_HR2_PstI_Fw | CGCGCGCTGCAG**TGGCTCTTGAAAGCCTTTGC** |
| ispA_HR2_XbaI_Rv | CGCGCGTCTAGA**CAACCAGATTGACATGAGCAAC** |
| menF_HR1_HindIII_Fw | TTGTTGAAGCTT**GCACTATATTGGTTGCTTCC** |
| menF_HR1_PstI_Rv | ATATATCTGCAG**CATAGAAATTATAACATAGCTCAC** |
| menF_HR2_PstI_Fw | TATATACTGCAG**GAAGCCTTATGACTTCTATTTAG** |
| menF_HR2_XbaI_Rv | ATATATTCTAGA**TTCAGAAGAATTGGCTCCTG** |

^a^In the primer names, the target gene, up or down stream homologous region (HR 1 or HR2), introduced restriction site, forward (Fw) or reverse (Rv) primer is indicated.

^b^In the primer sequences, the annealing sequences are bold, and introduced restriction sites are underlined.

**Table S2. Primers used for PCR amplifying the promoter and open reading frame for each target gene.**

| Name | Primer sequence 5’ – 3’^a^ |
| --- | --- |
| 45-menF-Fw | TTGAAAACCGCTACGGATCA**CATATAATTGTTGGCAATCG** |
| 45-menF-Rv | TCAAAGAAAGCTTGAGCTCT**GATATTCTAAATAGAAGTCATAAGG** |
| 45-0196_Fw | TTGAAAACCGCTACGGATCA**AGCGTTCTCGGGAGATTG** |
| 45-0196_Rv | TCAAAGAAAGCTTGAGCTCT**AACCTATCGGTGACAGGC** |
| 45-ispA_Fw | TCAAAGAAAGCTTGAGCTCT**CATTCTCTGTGAGAATTCTAAG** |
| 45-ispA_Rv | TTGAAAACCGCTACGGATCA**GAATTGGTGGCCAACAATATTC** |

^a^In the primer sequences, the annealing sequences are bold.

**Table S3. Primers used for PCR checking mutants with target gene knocked out.**

| Name | Primer sequence 5’ – 3’ |
| --- | --- |
| final check 0196 KO Fw | GCACACAATATTCAAAGTTGGC |
| final check 0196 KO Rv | CAAAGTCCACCAAATGAGTGC |
| final check ispA KO FW | GCATCACATCAGTAAATCCTCC |
| final check ispA KO Rv | GACGAGAAAGGAGAAGAAATTCC |
| final check menF KO Fw | GCAATCCCACGAACACTGAC |
| final check menF KO Rv | AGCATCATAGCTGTCCATGAC |

**Table S4. Quantity (Log LFQ intensity) of proteins involved in the primary metabolism in MG1363 and mutant under different cultivation conditions.** Values are averages from samples collected from 3 independent experiments, SEM values are shown below the average values. Detection limit in Log LFQ intensity: 6.7; Δ*0196* = Δ*llmg_0196.* Values in mutants that are significantly different (p < 0.05, fold change > 2) from strain MG1363 under the same cultivation conditions are highlighted in bold letters.

| **Protein** | **Gene** | **AN** | | | | **AE** | | | | **RES** | | | |
| --- | --- | --- | --- | --- | --- | --- | --- | --- | --- | --- | --- | --- | --- |
| **Function** | | **MG1363** | **Δ*menF*** | **Δ*ispA*** | **Δ*0196*** | **MG1363** | **Δ*menF*** | **Δ*ispA*** | **Δ*0196*** | **MG1363** | **Δ*menF*** | **Δ*ispA*** | **Δ*0196*** |
| A2RMN3 | *pflA* | 8.71 | 8.72 | 8.86 | 8.79 | 7.01 | ND | 7.03 | 7.04 | 7.70 | 6.97 | 7.37 | 6.98 |
| Pyruvate formate-lyase-activating enzyme | | 0.02 | 0.04 | 0.07 | 0.04 | 0.31 | - | 0.33 | 0.34 | 0.09 | 0.27 | 0.34 | 0.28 |
| O32799 | *pfl* | 10.26 | 10.26 | 10.43 | 10.25 | 9.60 | 9.65 | 9.86 | 9.59 | 9.93 | 9.69 | 9.91 | **9.56** |
| Formate acetyltransferase | | 0.03 | 0.02 | 0.02 | 0.00 | 0.02 | 0.02 | 0.03 | 0.01 | 0.04 | 0.01 | 0.01 | **0.01** |
| P0CI34 | *ldhB* | ND | ND | ND | ND | ND | ND | ND | ND | ND | ND | ND | ND |
| L-lactate dehydrogenase 2 | | - | - | - | - | - | - | - | - | - | - | - | - |
| A2RKA4 | *Ldh* | 10.71 | 10.66 | 10.60 | 10.68 | 10.53 | 10.56 | 10.37 | 10.52 | 10.45 | 10.55 | 10.43 | 10.45 |
| L-lactate dehydrogenase | | 0.03 | 0.01 | 0.02 | 0.02 | 0.04 | 0.01 | 0.02 | 0.03 | 0.01 | 0.04 | 0.02 | 0.02 |
| A2RL45 | *ldhX* | 8.21 | 8.34 | **8.76** | 8.24 | 7.33 | 7.69 | **8.53** | 7.70 | 7.69 | 7.86 | **8.64** | 7.08 |
| L-lactate dehydrogenase | | 0.13 | 0.07 | **0.04** | 0.08 | 0.32 | 0.01 | **0.03** | 0.03 | 0.01 | 0.06 | **0.02** | 0.38 |
| A2RHE1 | *pdhD* | 10.32 | 10.39 | 10.41 | 10.36 | 10.55 | 10.53 | 10.56 | 10.56 | 10.66 | 10.51 | 10.54 | 10.55 |
| Dihydrolipoyl dehydrogenase | | 0.01 | 0.02 | 0.01 | 0.03 | 0.01 | 0.04 | 0.01 | 0.01 | 0.02 | 0.01 | 0.00 | 0.02 |
| A2RHE2 | *pdhC* | 10.53 | 10.51 | 10.49 | 10.51 | 10.71 | 10.64 | 10.70 | 10.74 | 10.80 | 10.70 | 10.66 | 10.72 |
| Dihydrolipoamide acetyltransferase component of pyruvate dehydrogenase complex | | 0.02 | 0.04 | 0.04 | 0.04 | 0.02 | 0.02 | 0.04 | 0.03 | 0.01 | 0.01 | 0.02 | 0.03 |
| A2RHE3 | *pdhB* | 10.54 | 10.56 | 10.56 | 10.59 | 10.70 | 10.70 | 10.70 | 10.70 | 10.84 | 10.70 | 10.68 | 10.71 |
| Pyruvate dehydrogenase E1 component beta subunit | | 0.02 | 0.01 | 0.06 | 0.02 | 0.00 | 0.03 | 0.02 | 0.03 | 0.01 | 0.03 | 0.02 | 0.02 |
| A2RHE4 | *pdhA* | 10.23 | 10.19 | 10.20 | 10.19 | 10.37 | 10.35 | 10.34 | 10.37 | 10.54 | 10.40 | 10.36 | 10.41 |
| Pyruvate dehydrogenase E1 component alpha subunit | | 0.01 | 0.02 | 0.04 | 0.05 | 0.06 | 0.02 | 0.02 | 0.03 | 0.02 | 0.01 | 0.03 | 0.04 |
| A2RMM7 | *adhA* | 8.45 | 8.41 | 8.52 | 8.39 | 7.80 | 7.74 | 8.06 | 7.84 | 8.01 | 7.74 | 8.19 | 8.26 |
| Alcohol dehydrogenase, zinc-containing | | 0.02 | 0.04 | 0.03 | 0.04 | 0.02 | 0.12 | 0.11 | 0.04 | 0.04 | 0.06 | 0.03 | 0.08 |
| A2RNV2 | *adhE* | 10.66 | 10.68 | 10.71 | 10.67 | 9.48 | 9.54 | 9.42 | 9.47 | 8.51 | **9.54** | **9.29** | **9.01** |
| Aldehyde-alcohol dehydrogenase | | 0.03 | 0.02 | 0.02 | 0.02 | 0.02 | 0.03 | 0.02 | 0.02 | 0.03 | **0.01** | **0.01** | **0.02** |
| A2RJB5 | *pta* | 9.95 | 10.00 | 10.01 | 9.99 | 10.11 | 10.08 | 10.18 | 10.12 | 10.15 | 10.09 | 10.14 | 10.15 |
| Phosphate acetyltransferase | | 0.02 | 0.01 | 0.01 | 0.02 | 0.02 | 0.02 | 0.01 | 0.02 | 0.01 | 0.01 | 0.02 | 0.03 |
| A2RNG4 | *ackA2* | 9.69 | 9.71 | 9.70 | 9.69 | 9.71 | 9.70 | 9.75 | 9.71 | 9.82 | 9.74 | 9.71 | 9.70 |
| Acetate kinase | | 0.05 | 0.01 | 0.01 | 0.03 | 0.04 | 0.01 | 0.03 | 0.04 | 0.01 | 0.01 | 0.02 | 0.02 |
| A2RNG5 | *ackA1* | 9.97 | 9.96 | 9.95 | 9.97 | 9.93 | 9.96 | 9.86 | 9.90 | 9.82 | 9.94 | 9.88 | 9.87 |
| Acetate kinase | | 0.05 | 0.04 | 0.01 | 0.00 | 0.01 | 0.03 | 0.02 | 0.03 | 0.02 | 0.01 | 0.03 | 0.01 |
| A2RKT5 | *als* | 9.81 | 9.82 | 9.83 | 9.77 | 10.18 | 10.09 | 10.19 | 10.13 | 10.38 | 10.13 | 10.21 | 10.23 |
| Acetolactate synthase large subunit | | 0.00 | 0.03 | 0.01 | 0.01 | 0.01 | 0.02 | 0.02 | 0.02 | 0.03 | 0.01 | 0.01 | 0.02 |
| A2RKQ8 | *ilvB* | ND | 7.12 | 7.49 | 7.11 | 7.04 | 7.03 | 7.81 | 7.03 | 7.09 | 7.72 | 7.92 | 7.35 |
| Acetolactate synthase | | - | 0.42 | 0.40 | 0.41 | 0.34 | 0.33 | 0.05 | 0.33 | 0.39 | 0.05 | 0.06 | 0.33 |
| P77880 | *aldB* | 8.83 | 8.74 | 8.67 | 8.70 | 8.79 | 8.86 | 8.95 | 8.83 | 8.86 | 8.87 | 8.72 | 8.84 |
| Alpha-acetolactate decarboxylase | | 0.02 | 0.03 | 0.03 | 0.07 | 0.03 | 0.07 | 0.04 | 0.11 | 0.03 | 0.04 | 0.09 | 0.04 |
| A2RLP5 | *butA* | 9.18 | 9.18 | 9.45 | 9.17 | 9.21 | 9.19 | 9.38 | 9.26 | 9.59 | **9.21** | 9.46 | 9.47 |
| Acetoin reductase | | 0.01 | 0.01 | 0.02 | 0.03 | 0.03 | 0.01 | 0.01 | 0.02 | 0.01 | **0.05** | 0.03 | 0.03 |

**Table S5. Quantity (Log LFQ intensity) of proteins that showed significant changes across the strains or cultivation conditions (a non-exhaustive list).** Values are averages from samples collected from 3 independent experiments, SEM values are shown below the average values. Detection limit in Log LFQ intensity: 6.7; Δ*0196* = Δ*llmg_0196.* Values in mutants that are significantly different (p < 0.05, fold change > 2) from strain MG1363 under the same cultivation conditions are highlighted in bold letters.

| **Protein** | **Gene** | **AN** | | | | **AE** | | | | **RES** | | | |
| --- | --- | --- | --- | --- | --- | --- | --- | --- | --- | --- | --- | --- | --- |
| **Function** | | **MG1363** | **Δ*menF*** | **Δ*ispA*** | **Δ*0196*** | **MG1363** | **Δ*menF*** | **Δ*ispA*** | **Δ*0196*** | **MG1363** | **Δ*menF*** | **Δ*ispA*** | **Δ*0196*** |
| A2RKD8 | *pabB* | 8.33 | 7.33 | 8.48 | 8.54 | ND | 7.26 | 7.69 | ND | ND | 7.27 | 7.25 | ND |
| Aminodeoxychorismate synthase | | 0.03 | 0.63 | 0.07 | 0.05 | - | 0.56 | 0.49 | - | - | 0.57 | 0.55 | - |
| A2RK13 | *llmg_1028* | 8.88 | 8.76 | 8.98 | 8.78 | 8.49 | 8.37 | 8.70 | 8.34 | 7.58 | **8.31** | **8.75** | **8.40** |
| Putative NAD(P)H nitroreductase | | 0.03 | 0.01 | 0.03 | 0.07 | 0.05 | 0.08 | 0.05 | 0.11 | 0.14 | **0.04** | **0.03** | **0.03** |
| A2RIQ1 | *dfpA* | 8.87 | 8.72 | 8.76 | 8.68 | 8.50 | 8.49 | 8.29 | 8.42 | 8.30 | 8.28 | 8.22 | 7.89 |
| Putative DNA/pantothenate metabolism flavoprotein | | 0.05 | 0.07 | 0.03 | 0.06 | 0.04 | 0.07 | 0.08 | 0.07 | 0.11 | 0.08 | 0.09 | 0.23 |
| A2RLD1 | *llmg_1522* | 7.92 | 7.76 | 7.63 | 7.75 | 7.78 | 7.70 | 7.77 | 7.81 | 7.57 | 7.75 | 7.67 | 7.43 |
| Putative membrane protein | | 0.10 | 0.08 | 0.01 | 0.04 | 0.09 | 0.04 | 0.03 | 0.06 | 0.06 | 0.02 | 0.09 | 0.37 |
| A2RIP8 | *yidC* | 9.45 | 9.34 | 9.29 | 9.32 | 9.32 | 9.26 | 9.21 | 9.25 | 9.10 | 9.24 | 9.17 | 9.23 |
| Membrane protein insertase | | 0.04 | 0.06 | 0.03 | 0.04 | 0.04 | 0.03 | 0.11 | 0.10 | 0.07 | 0.02 | 0.01 | 0.02 |
| A2RNY5 | *recX* | 8.28 | 8.10 | 8.27 | 8.28 | 7.82 | 7.85 | **8.17** | 7.77 | 7.95 | 8.01 | 8.15 | 8.04 |
| Regulatory protein | | 0.08 | 0.08 | 0.04 | 0.05 | 0.09 | 0.06 | **0.04** | 0.06 | 0.03 | 0.04 | 0.10 | 0.08 |
| A2RHL7 | *llmg_0146* | 9.24 | 9.26 | 9.38 | 9.30 | 9.24 | 9.32 | 9.36 | 9.32 | 9.55 | 9.32 | 9.42 | 9.33 |
| Aryl-alcohol dehydrogenase | | 0.06 | 0.01 | 0.04 | 0.09 | 0.04 | 0.02 | 0.04 | 0.07 | 0.04 | 0.02 | 0.03 | 0.04 |
| Q9S5Z2 | *clpE* | 10.28 | 10.25 | 10.33 | 10.21 | 10.29 | 10.28 | 10.31 | 10.27 | 10.59 | 10.32 | 10.28 | 10.39 |
| ATP-dependent Clp protease ATP-binding subunit | | 0.02 | 0.01 | 0.02 | 0.01 | 0.02 | 0.01 | 0.03 | 0.01 | 0.01 | 0.02 | 0.02 | 0.01 |
| A2RMQ7 | *llmg_2023* | 10.02 | 10.08 | 10.04 | 10.11 | 10.10 | 10.07 | 10.05 | 10.13 | 10.33 | 10.13 | 10.09 | 10.21 |
| Universal stress protein A | | 0.01 | 0.03 | 0.06 | 0.03 | 0.04 | 0.04 | 0.03 | 0.01 | 0.04 | 0.02 | 0.02 | 0.02 |
| A2RLA1 | *mntH* | 8.53 | 8.47 | 8.46 | 8.53 | 8.93 | 8.83 | 8.69 | 8.86 | 8.87 | 8.88 | 8.65 | 8.96 |
| Divalent metal cation transporter | | 0.05 | 0.09 | 0.08 | 0.09 | 0.07 | 0.04 | 0.09 | 0.03 | 0.03 | 0.06 | 0.06 | 0.10 |
| A2RJM2 | *llmg_0880* | 9.17 | 9.21 | 9.20 | 9.19 | 9.17 | 9.15 | 9.10 | 9.16 | 9.52 | **9.11** | **9.14** | 9.26 |
| Putative oxidoreductase | | 0.05 | 0.06 | 0.04 | 0.07 | 0.03 | 0.02 | 0.02 | 0.04 | 0.03 | **0.03** | **0.02** | 0.02 |
| A2RM94 | *qor* | 8.57 | 8.72 | 8.54 | 8.64 | 8.67 | 8.68 | 8.62 | 8.68 | 8.92 | 8.78 | **8.55** | 8.66 |
| Zinc-type alcohol dehydrogenase-like protein | | 0.01 | 0.04 | 0.01 | 0.07 | 0.02 | 0.10 | 0.05 | 0.06 | 0.03 | 0.04 | **0.02** | 0.04 |
| A2RHS0 | *msrB* | 8.86 | 8.75 | 8.82 | 8.82 | 8.89 | 8.89 | 8.88 | 8.84 | 9.21 | **8.91** | **8.89** | 8.95 |
| Peptide methionine sulfoxide reductase | | 0.00 | 0.03 | 0.04 | 0.05 | 0.02 | 0.01 | 0.04 | 0.05 | 0.02 | **0.01** | **0.04** | 0.04 |
| A2RI61 | *fhuD* | 8.81 | 8.73 | 8.73 | 8.79 | 9.30 | 9.25 | 9.34 | 9.31 | 9.17 | 9.16 | 9.33 | 9.20 |
| Ferrichrome ABC transporter substrate binding protein | | 0.03 | 0.04 | 0.06 | 0.03 | 0.01 | 0.02 | 0.04 | 0.04 | 0.02 | 0.04 | 0.02 | 0.03 |
| A2RJT8 | *msrA* | 9.11 | 9.08 | 9.14 | 9.05 | 9.20 | 9.30 | 9.16 | 9.18 | 9.50 | **9.16** | **9.15** | 9.32 |
| Peptide methionine sulfoxide reductase | | 0.07 | 0.01 | 0.01 | 0.06 | 0.06 | 0.03 | 0.02 | 0.03 | 0.02 | **0.03** | **0.02** | 0.05 |
| A2RKC2 | *mtsA* | 9.40 | 9.36 | 9.36 | 9.44 | 9.63 | 9.64 | 9.50 | 9.63 | 9.82 | 9.62 | **9.50** | 9.70 |
| Manganese ABC transporter substrate binding protein | | 0.03 | 0.03 | 0.03 | 0.03 | 0.05 | 0.06 | 0.05 | 0.05 | 0.02 | 0.04 | **0.03** | 0.03 |
| A2RK08 | *fur* | 8.30 | 8.38 | 8.27 | 8.44 | 8.38 | 8.47 | 8.30 | 8.38 | 8.75 | 8.52 | **8.44** | 8.48 |
| Ferric uptake regulation protein | | 0.12 | 0.03 | 0.03 | 0.06 | 0.05 | 0.01 | 0.15 | 0.13 | 0.04 | 0.03 | **0.04** | 0.14 |
| A2RM21 | *metC* | 9.08 | 9.06 | 8.92 | 9.06 | 9.42 | 9.36 | 9.17 | 9.37 | 9.56 | **9.26** | **9.14** | 9.33 |
| Cystathionine beta-lyase | | 0.02 | 0.04 | 0.03 | 0.04 | 0.04 | 0.03 | 0.04 | 0.02 | 0.04 | **0.01** | **0.02** | 0.00 |
| A2RNK4 | *llmg_2331* | 9.26 | 9.18 | 9.18 | 9.17 | 9.32 | 9.30 | 9.26 | 9.30 | 9.76 | **9.27** | **9.23** | **9.44** |
| Pyridine nucleotide-disulfide oxidoreductase | | 0.00 | 0.02 | 0.04 | 0.02 | 0.02 | 0.01 | 0.05 | 0.03 | 0.01 | **0.01** | **0.02** | **0.04** |
| A2RKC0 | *mtsB* | 9.12 | 9.02 | 9.11 | 9.18 | 9.42 | 9.40 | 9.39 | 9.43 | 9.65 | 9.43 | 9.36 | 9.49 |
| Manganese ABC transporter ATP binding protein | | 0.03 | 0.02 | 0.02 | 0.02 | 0.03 | 0.03 | 0.02 | 0.08 | 0.02 | 0.04 | 0.03 | 0.04 |
| A2RK64 | *uspA2* | 9.57 | 9.54 | 9.69 | 9.49 | 9.87 | 9.76 | 9.89 | 9.89 | 10.12 | 9.83 | 9.90 | 10.09 |
| Universal stress protein A2 | | 0.01 | 0.04 | 0.01 | 0.05 | 0.01 | 0.02 | 0.05 | 0.04 | 0.02 | 0.02 | 0.01 | 0.06 |
| A2RJC6 | *llmg_0776* | 8.59 | 8.45 | 8.41 | 8.59 | 9.06 | 8.93 | 9.01 | 8.98 | 9.16 | 8.98 | 8.97 | 9.06 |
| Ferredoxin--NADP reductase | | 0.05 | 0.01 | 0.04 | 0.03 | 0.03 | 0.02 | 0.02 | 0.06 | 0.02 | 0.02 | 0.04 | 0.04 |
| A2RLR6 | *uspA* | 9.40 | 9.38 | 9.49 | 9.40 | 9.53 | 9.54 | 9.58 | 9.52 | 10.02 | **9.50** | **9.64** | 9.78 |
| Universal stress protein | | 0.04 | 0.06 | 0.03 | 0.03 | 0.01 | 0.02 | 0.06 | 0.03 | 0.04 | **0.04** | **0.06** | 0.12 |
| A2RHF0 | *osmC* | 8.93 | 9.06 | 8.89 | 8.97 | 9.19 | 9.14 | 9.14 | 9.23 | 9.60 | **9.23** | **9.11** | 9.28 |
| Osmotically inducible protein | | 0.14 | 0.02 | 0.14 | 0.04 | 0.05 | 0.05 | 0.06 | 0.12 | 0.02 | **0.06** | **0.03** | 0.12 |
| A2RHZ1 | *llmg_0276* | 10.44 | 10.46 | 10.55 | 10.47 | 10.61 | 10.60 | 10.58 | 10.57 | 11.18 | **10.65** | **10.62** | **10.77** |
| Oxidoreductase, aldo/keto reductase | | 0.06 | 0.04 | 0.03 | 0.04 | 0.02 | 0.03 | 0.02 | 0.03 | 0.03 | 0.03 | 0.03 | 0.02 |
| A2RHG2 | *cysD* | 8.03 | 8.28 | **8.37** | 8.14 | 8.63 | 8.53 | 8.66 | 8.69 | 8.81 | 8.60 | 8.70 | 8.51 |
| O-acetylhomoserine sulfhydrylase | | 0.08 | 0.10 | **0.03** | 0.11 | 0.03 | 0.03 | 0.05 | 0.02 | 0.03 | 0.01 | 0.06 | 0.06 |
| A2RI58 | *fhuC* | 7.05 | 7.80 | 7.86 | 7.41 | 8.04 | 7.88 | **8.41** | 8.12 | 8.11 | 7.93 | 8.27 | 8.05 |
| Ferrichrome ABC transporter | | 0.35 | 0.10 | 0.00 | 0.35 | 0.03 | 0.03 | **0.10** | 0.08 | 0.08 | 0.04 | 0.04 | 0.11 |
| A2RNJ4 | *poxL* | ND | ND | 6.97 | ND | ND | ND | ND | ND | 8.17 | **ND** | **ND** | **ND** |
| Pyruvate oxidase | | - | - | 0.27 | - | - | - | - | - | 0.08 | **-** | **-** | **-** |
| A2RMS9 | *nifS* | 8.89 | 8.91 | 8.95 | 8.92 | 8.70 | 8.64 | **9.06** | 8.68 | 9.02 | 8.93 | 9.09 | 8.97 |
| Putative iron-sulfur cofactor synthesis protein | | 0.10 | 0.01 | 0.04 | 0.05 | 0.05 | 0.05 | **0.05** | 0.08 | 0.03 | 0.04 | 0.01 | 0.00 |
| P42370 | *hrcA* | 8.73 | 8.68 | 9.01 | 8.66 | 8.18 | 8.19 | **8.91** | 8.24 | 8.53 | **8.08** | 8.80 | 8.38 |
| Heat-inducible transcription repressor | | 0.02 | 0.02 | 0.04 | 0.01 | 0.06 | 0.02 | **0.03** | 0.04 | 0.04 | **0.05** | 0.01 | 0.03 |
| A2RLX6 | *merP* | 7.76 | 7.76 | 7.62 | 7.74 | 7.16 | 7.46 | **7.87** | 7.43 | 8.09 | 7.68 | 7.84 | 7.80 |
| Mercuric reductase | | 0.15 | 0.03 | 0.21 | 0.09 | 0.23 | 0.15 | **0.08** | 0.07 | 0.16 | 0.05 | 0.09 | 0.11 |
| A2RHR8 | *feoB* | 7.87 | 7.86 | 7.80 | 8.06 | 6.94 | ND | 7.50 | ND | ND | 6.97 | **7.68** | 6.92 |
| Ferrous iron transport protein B | | 0.06 | 0.08 | 0.04 | 0.09 | 0.24 | - | 0.08 | - | - | 0.27 | **0.04** | 0.22 |
| A2RNH7 | *dpsA* | 9.92 | 9.87 | 9.88 | 9.90 | 9.08 | 8.94 | 9.21 | 9.11 | 9.21 | 9.08 | 9.25 | 8.98 |
| Non-heme iron-binding ferritin | | 0.31 | 0.31 | 0.34 | 0.30 | 0.07 | 0.07 | 0.05 | 0.05 | 0.03 | 0.11 | 0.18 | 0.07 |
| A2RMF4 | *llmg_1915* | 9.32 | 9.26 | 9.31 | 9.24 | 8.73 | 8.57 | 8.88 | 8.67 | 8.95 | **8.57** | 8.87 | 8.73 |
| Putative Fe-S oxidoreductase | | 0.06 | 0.02 | 0.04 | 0.04 | 0.04 | 0.06 | 0.05 | 0.07 | 0.07 | **0.04** | 0.03 | 0.07 |
| A2RHR9 | *feoA* | 7.07 | 7.75 | 7.40 | 7.71 | 7.07 | 7.79 | 7.36 | 7.16 | 7.12 | ND | 7.72 | ND |
| Ferrous iron transport protein A | | 0.37 | 0.01 | 0.35 | 0.06 | 0.37 | 0.19 | 0.33 | 0.46 | 0.42 | - | 0.01 | - |
| A2RJ14 | *llmg_0661* | 7.33 | 7.53 | 7.26 | 7.28 | 7.60 | 7.30 | 7.55 | 7.59 | 7.54 | **ND** | 7.26 | 7.28 |
| CorA like magnesium and cobalt transport protein | | 0.32 | 0.03 | 0.28 | 0.29 | 0.06 | 0.30 | 0.03 | 0.03 | 0.06 | **-** | 0.28 | 0.29 |
| A2RI68 | *ahpC* | 10.47 | 10.53 | 10.42 | 10.44 | 10.43 | 10.40 | 10.31 | 10.44 | 10.18 | 10.42 | 10.23 | 10.29 |
| Alkyl hydroperoxide reductase subunit C | | 0.04 | 0.01 | 0.01 | 0.05 | 0.01 | 0.05 | 0.04 | 0.02 | 0.02 | 0.03 | 0.01 | 0.02 |
| A2RI69 | *ahpF* | 10.22 | 10.24 | 10.19 | 10.26 | 10.22 | 10.17 | 10.06 | 10.20 | 9.94 | 10.20 | 10.01 | 10.07 |
| Alkyl hydroperoxide reductase subunit F | | 0.02 | 0.02 | 0.02 | 0.01 | 0.01 | 0.01 | 0.03 | 0.01 | 0.04 | 0.02 | 0.01 | 0.03 |
| A2RLU6 | *copB* | 8.64 | 8.53 | 8.44 | 8.59 | 8.58 | 8.49 | 8.52 | 8.56 | 8.66 | 8.47 | 8.41 | 8.56 |
| Copper-potassium transporting ATPase B | | 0.03 | 0.02 | 0.05 | 0.04 | 0.03 | 0.04 | 0.07 | 0.01 | 0.03 | 0.02 | 0.06 | 0.03 |
| A2RLX5 | *copA* | 8.62 | 8.70 | 8.53 | 8.64 | 8.24 | 8.29 | 8.24 | 8.23 | 8.25 | 8.30 | **8.60** | 8.37 |
| Copper/potassium-transporting ATPase | | 0.12 | 0.07 | 0.08 | 0.07 | 0.04 | 0.01 | 0.16 | 0.03 | 0.06 | 0.07 | **0.09** | 0.03 |
| A2RNE6 | *cutC* | 9.66 | 9.57 | 9.54 | 9.59 | 9.57 | 9.52 | 9.52 | 9.56 | 9.58 | 9.52 | 9.50 | 9.56 |
| Copper homeostasis protein | | 0.01 | 0.02 | 0.04 | 0.04 | 0.02 | 0.04 | 0.06 | 0.02 | 0.03 | 0.03 | 0.01 | 0.01 |
